# Mechanism of high energy efficiency of carbon fixation by sulfur-oxidizing symbionts revealed by single-cell analyses and metabolic modeling

**DOI:** 10.1101/2023.11.25.568684

**Authors:** M. Kleiner, L. Polerecky, C. Lott, C. Bergin, S. Häusler, M. Liebeke, C. Wentrup, N. Musat, M. M. M. Kuypers, N. Dubilier

**Affiliations:** Max Planck Institute for Marine Microbiology, Celsiusstr. 1, 28359 Bremen, Germany; Department of Plant and Microbial Biology, North Carolina State University, Raleigh, NC, 27695, USA; Department of Earth Sciences, Faculty of Geosciences, Utrecht University, The Netherlands; HYDRA Institute for Marine Sciences, Elba Field Station, Via del Forno 80, Località, Fetovaia, 57034 Campo nell’Elba (LI), Italy; Department of Cell and Molecular Biology, Uppsala University, and Microbial Single Cell Genomics unit, Science for Life Laboratory, Uppsala, Sweden; Division of Metabolomics, Institute of Human Nutrition and Food Science, University of Kiel, Kiel, Germany; Biokar Diagnostics Groupe Solabia, France; Department of Biology, Aarhus University, Aarhus, Denmark

**Keywords:** symbiosis, Oligochaeta, *Olavius algarvensis*, chemolithoautotroph, sulfur oxidizer, carbon fixation, single cell, nanoSIMS, Calvin cycle, energy efficiency

## Abstract

In chemosynthetic symbioses between marine invertebrates and autotrophic sulfur-oxidizing bacteria the symbionts feed their host by producing organic compounds from CO_2_ using reduced sulfur compounds as an energy source. One such symbiosis, the gutless marine *worm Olavius algarvensis* harbors at least five bacterial symbionts of which four have the genetic potential for an autotrophic metabolism.

In this study we combined single-cell analyses of CO_2_ fixation, CO_2_ release and bulk uptake, with measurements of O_2_ respiration, sulfur content, and polyhydroxyalkanoate content, as well as mathematical modelling to investigate how energy derived from sulfur oxidation drives carbon fluxes within the symbiosis and between the holobiont and its habitat. We found that under aerobic conditions without external energy sources only the primary symbiont, *Ca*. Thiosymbion algarvensis, fixed carbon. This symbiont relied on internal sulfur storage for energy production. Our model showed that the apparent efficiency of carbon fixation driven by sulfur oxidation in the symbiosis was higher than thermodynamically feasible if only stored sulfur was considered as source of energy and reducing equivalents. The model and additional calculations showed that reducing equivalents must be derived from a different source than energy. We identified the large amounts of polyhdroxyalkanoate stored by the symbiont as the likely source of reducing equivalents for carbon fixation in the symbiont which boosts the yield of sulfur-driven carbon fixation. The model also showed that heterotrophic carbon fixation by host tissue is not negligible and has to be considered when assessing transfer of carbon from the symbionts to the host.

## Introduction

Chemosynthetic symbioses between marine invertebrates and chemolithoautotrophic sulfur-oxidizing bacteria (SOX) are found in a broad range of animal phyla and marine environments (Dubilier et al., 2008; Sogin et al., 2021). In these symbioses the symbionts “feed” their host by producing organic compounds from CO_2_ using reduced sulfur compounds as an energy source (Cavanaugh et al., 2013). The SOX symbionts are often Gammaproteobacteria that employ the Calvin-Benson-Bassham (CBB) cycle for autotrophic carbon fixation (Dubilier et al., 2008; Kleiner et al., 2012a). A recent study discovered that sulfur-oxidation driven carbon fixation is much more efficient in these symbiotic SOX as compared to free-living SOX (Klatt and Polerecky, 2015). Klatt and Polerecky (2015) speculated that potentially competition of the symbiotic SOX with their host animal for oxygen may select for higher efficiency, however, a mechanistic understanding of how the higher efficiency is achieved is currently lacking.

Gutless marine, phallodrilines (oliogochaetes, Annelida, Clitellata, Naididae *sensu* (Erseus et al., 2008)) are commonly found in tropical and subtropical sediments. They live in symbiosis with a phylogenetically and metabolically diverse community of bacterial endosymbionts. These worms lack a mouth, intestinal tract and excretory organs and depend on their symbionts that provide them with nutrition and recycle their waste compounds (Dubilier et al., 2006; Woyke et al., 2006; Kleiner et al., 2012b). All gutless oligochaetes harbor a primary, gammaproteobacterial symbiont of the genus *Candidatus* Thiosymbion (previously Gamma1 or γ1 symbiont) (Zimmermann et al., 2016), with the exception of *Inanidrilus exumae* in which *Ca*. Thiosymbion has been replaced by another Gammaproteobacterium (Bergin et al., 2018). *Ca.* Thiosymbion stores large amounts of sulfur (Krieger et al., 2000) and the carbon storage compound polyhydroxyalkanoate (PHA) (Giere et al., 1988), and uses reduced sulfur compounds as an energy source for the autotrophic fixation of CO_2_ (Dubilier et al., 2001; Dubilier et al., 2006). In addition to their primary symbionts, all hosts are associated with a phylogenetically diverse assemblage of secondary symbionts (Dubilier et al., 2006; Mankowski et al., 2021). The widespread dominance of the primary chemolithoautotrophic, sulfur-oxidizing *Ca*. Thiosymbion in all gutless oligochaete species has led to the assumption that it plays a key role in the nutrition of these hosts.

One of the most well-studied gutless oligochaetes is *Olavius algarvensis* from the Mediterranean. This species harbors two sulfur-oxidizing symbionts (*Ca*. Thiosymbion algarvensis and the γ3-symbiont), two sulfate-reducing deltaproteobacterial symbionts (the δ1 and δ4 symbionts), and a spirochaete (Ruehland et al., 2008; Sato et al., 2022). Metagenomic, metaproteomic, metabolomic and physiological analyses of the *O. algarvensis* symbiont community revealed numerous metabolic pathways and interactions among the symbionts as well as between these and their host (Dubilier et al., 2001; Woyke et al., 2006; Kleiner et al., 2012b; Kleiner et al., 2015). These include the syntrophic exchange of oxidized and reduced sulfur compounds between the sulfur-oxidizing symbionts and the sulfate-reducing δ symbionts, as well as pathways for autotrophic CO_2_ fixation in both the sulfur-oxidizing and the sulfate-reducing symbionts. The syntrophic sulfur compound exchange is enabled by worm migration between the oxic and anoxic layers of the sediment allowing temporally separated sulfate reduction and sulfur oxidation to occur (Woyke et al., 2006). Additionally, *Ca.* T. algarvensis was shown to have the capability to assimilate organic host-waste products into PHA using a novel metabolic pathway involving elements of the 3-hydroxypropionate bicycle (Kleiner et al., 2012b).

Given that *O. algarvensis* is dependent on its symbionts for its nutrition, one of the key questions is how nutrition is transferred to the host. A recent transcriptomic and proteomic study provided strong, but indirect evidence that the host “feeds” on the symbionts by endocytosing them (Wippler et al., 2016). Direct evidence for transfer of nutrients from symbiont to the host has so far been lacking.

To address the question of how high efficiencies are achieved by symbiotic SOX and how carbon flows through a chemosynthetic symbiosis we followed the flow of labelled ^13^C in the *O. algarvensis* symbiosis, focusing on the autotrophic activity of *Ca.* Thiosymbion. By incubating worms in the absence of external electron donors and under oxic conditions only *Ca.* Thiosymbion could fix inorganic carbon. Under these conditions, we expected, based on our previous analyses, that only *Ca.* Thiosymbion would have a source of energy, namely their internal stores of sulfur. In contrast, the γ3 symbionts do not appear to be able to store sulfur (Woyke et al., 2006). The oxic conditions further ensured that only *Ca.* Thiosymbion would be active, because the γ3 symbionts and the sulfate-reducing δ symbionts are not able to use oxygen as a terminal electron acceptor (Kleiner et al., 2012b). We analyzed the carbon in multiple compartments of the symbiosis to quantify the fluxes. Using isotopically labelled bicarbonate, we examined the incorporation of inorganic carbon in bulk analyses of whole worms as well as in single cells. For the latter, we combined nanometer-scale secondary ion mass spectrometry (nanoSIMS) with specific halogen *in situ* hybridization (HISH) (Musat et al., 2008; Musat et al., 2016) to distinguish *Ca.* Thiosymbion from the four other co-occurring symbionts. Additionally, we measured oxygen respiration rates and the sulfur content of worms, as well as the PHA content of purified *Ca*. Thiosymbion cells. Finally, we developed a mathematical model to quantify the rates and efficiency of inorganic carbon uptake by *Ca.* Thiosymbion and the transfer of organic carbon to host tissues.

## Results

### Model construction and assumptions

We constructed a model of carbon and energy flow in the *O. algarvensis* symbiosis assuming oxic conditions without external e^-^-donors and oxygen as the sole e^-^-acceptors (Fig. 1). These conditions are the ones met by the worm when it shuttles to the oxic layer of the sediment. We implemented the model mathematically in Matlab to (i) quantify carbon fluxes within the symbiosis and between the symbiosis and its environment, (ii) to quantify CO_2_ fixation rates of the symbionts, (iii) to gain insights into how fast CO_2_ fixed by the symbionts is transferred to the worm and (iv) to elucidate the role of the abundant storage compounds PHA and sulfur in CO_2_ fixation and carbon fluxes.

**Fig. 1:**
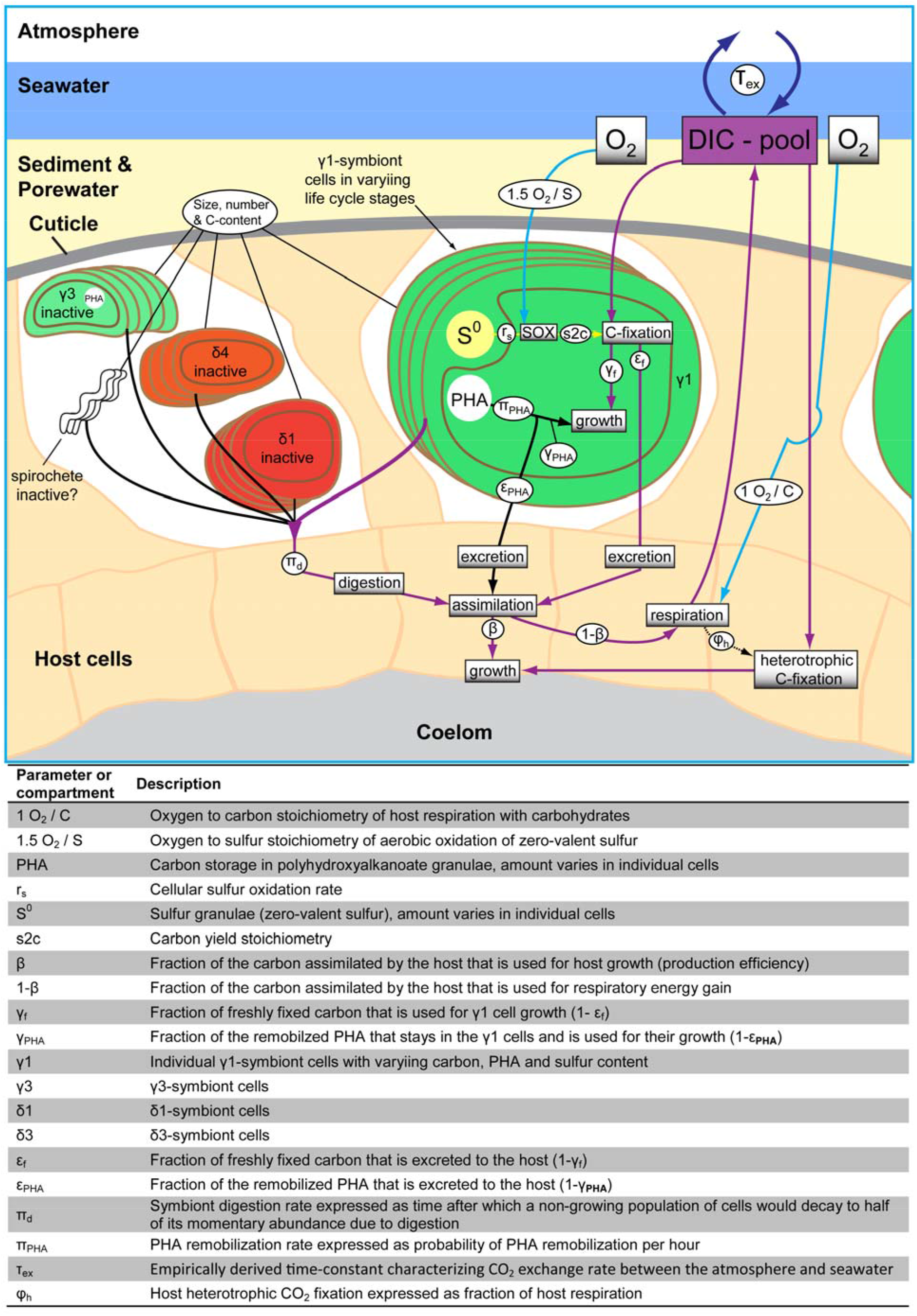
Conceptual model of carbon flow in *O. algarvensis*. The role of sulfur oxidation and oxygen respiration in carbon fluxes is included. Purple arrows indicate flow of freshly fixed carbon, black arrows indicate flow of pre-existing/stored carbon, and blue arrows indicate role of oxygen.

The two major compartments in the model are (1) the seawater outside of the worm and (2) the whole worm symbiosis containing six sub compartments – the five symbionts (*Ca*. Thiosymbion, γ3, δ1, δ4 and the spirochaete) and the host tissue. As a starting condition for the model we simulated worms freshly taken from the sediment. The *Ca*. Thiosymbion cells in such worms are usually filled with large amounts of stored sulfur (S^0^) and PHA. In the absence of external e^-^-donors, stored sulfur is thought to be the only energy source for CO_2_ fixation by the symbionts. PHA, which is generated from host waste products under anoxic conditions, is thought to be remobilized under oxic conditions (Kleiner et al., 2012b). A detailed overview of all literature-based assumptions for model construction can be found in the Supplementary Methods (SI METHODS), and the data used to fit model parameters can be found in the results below and in Table S1. The initial conditions file in the final model (see supplemental .zip folder) gives all model parameters, their estimated value and explanations of how values were obtained.

As the basis for fitting our model with the experimental data we defined an “average worm” based on the averages from our worm carbon bulk measurements (see below). An average worm thus has a carbon content of 5 μmol (60 μg C) and a biovolume of 0.532 µl. Wherever possible we used carbon content measures or biovolumes to convert rates and concentrations of different measurements to be comparable to the “average worm” (e. g. sulfur content and respiration rates).

### NanoSIMS: Only *Ca*. Thiosymbion fixes carbon under aerobic conditions without external energy source

Combined nanoSIMS and catalyzed reporter deposition (CARD) halogen *in situ* hybridization (HISH) (Musat et al., 2008) analyses allowed the clear identification of four of the five symbionts based on their size and hybridization signal with the general gammaproteobacterial probe (GAM42a)(Fig. 2). The spirochaete symbionts, which are estimated to account for at most 10% of the *O. algarvensis* symbiotic community (Ruehland et al., 2008), were not observed by nanoSIMS.

**Fig. 2:**
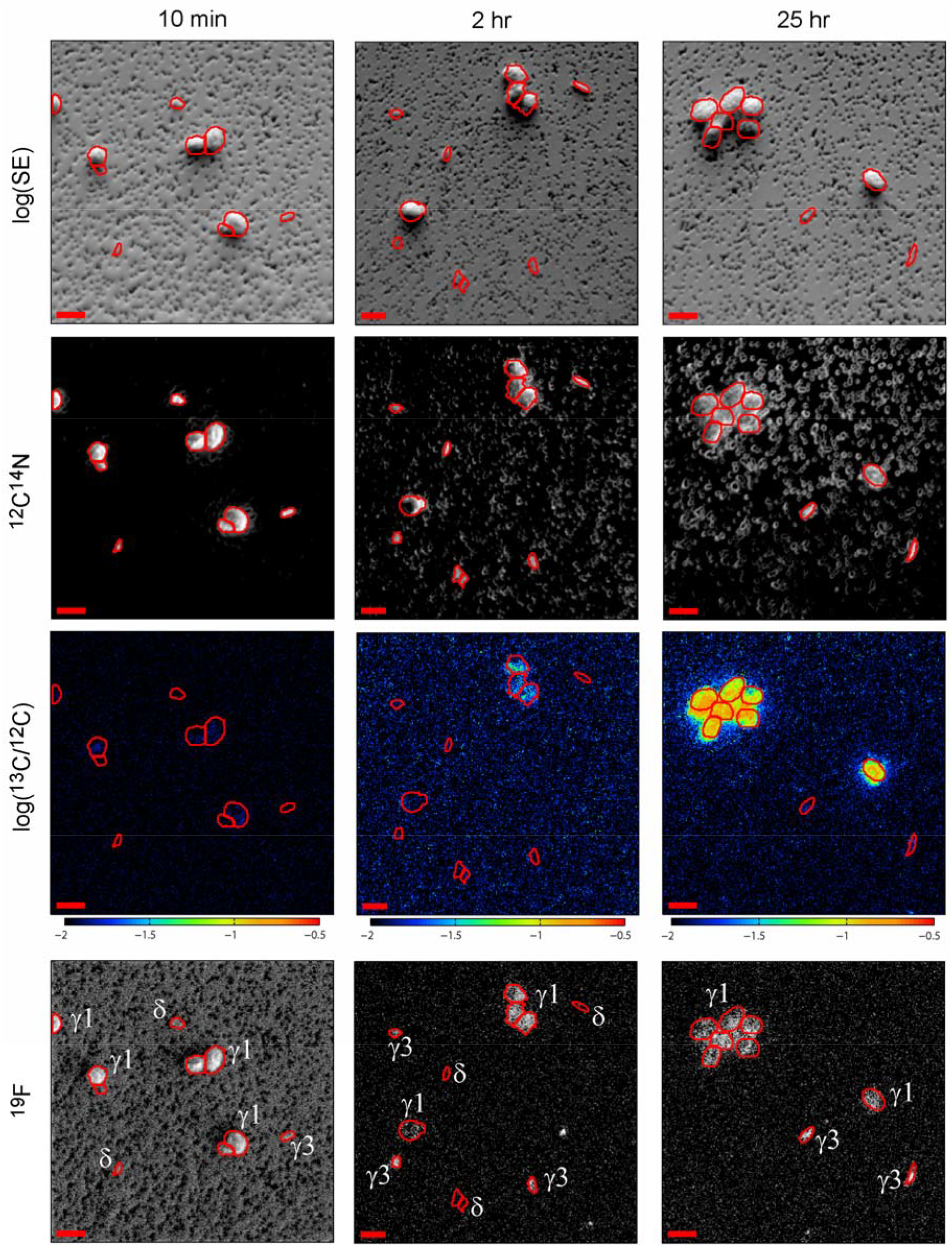
NanoSIMS images of individual symbiont cells at different incubation time points. Top row, secondary electron (SE) images. Cell outlines were drawn based on the SE and ^12^C^14^N images. Species identity was assigned based on the ^19^F signal of CARD-HISH hybridization with the gammaproteobacteria specific probe GAM42a and cell size. Scale bars are 2 µm.

Analyses of ^13^C/^12^C ratio images of the *O. algarvensis* symbionts revealed that inorganic carbon was only fixed by *Ca*. Thiosymbion (Fig. 2), as predicted. ^13^C abundance in *Ca.* Thiosymbion cells increased from 0.0109 (i.e. natural abundance; SE=0.00006, N=17) at time zero to 0.0124 (SE=0.0001, N=55) after the first 10 minutes of incubation, and further increased to 0.09 (SE=0.003, N=40) after 25 hours (Fig. 3, circles). In contrast, no significant increase in ^13^C abundance above natural levels was observed in the γ3 and δ symbionts even after 25 hours of incubation (Fig. 3, crosses; 0.0110, SE=0.0001, N=18; ANOVA, p=0.93).

**Fig. 3:**
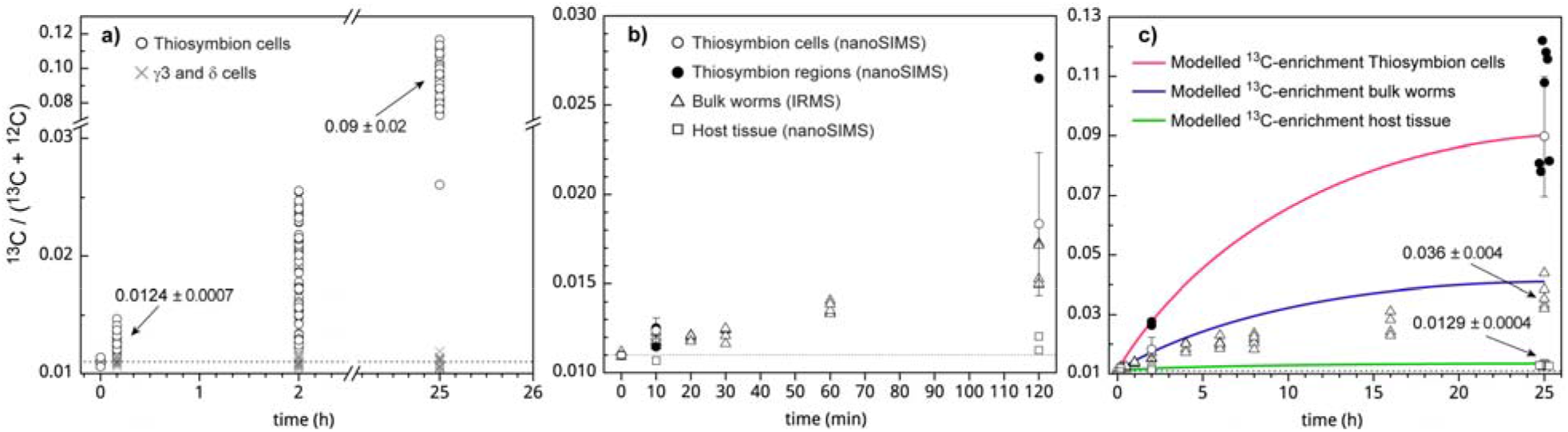
Measured and modelled ^13^C-enrichment in Olavius symbiosis compartments. **a)** Enrichment of ^13^C in single symbiont cells measured with nanoSIMS. Individual data points correspond to single symbiont cells. Note the break in the x- and y-axes in graph. **b)** ^13^C-enrichment in the first two hours of the experiment in whole worms, regions in worm cross-sections and single *Cand.* Thiosymbion cells. Averages for three pooled worms (triangles), and averages over the *Cand.* Thiosymbion and host tissue regions in single worm sections (filled circles and squares). **c)** Experimental data in comparison with the modelled data (fitted model). The legend for the experimental data is the same as in b). For single *Cand.* Thiosymbion cells the average and standard deviation are shown. Dotted horizontal line represents the natural ^13^C-abundance. Averages for selected data points are shown for orientation in a) and c).

### NanoSIMS: ^13^C enrichment in host tissue already observed after 2 hours

NanoSIMS measurements of tissue sections from worms incubated for 25 hours revealed strong ^13^C enrichment in regions containing *Ca*. Thiosymbion, as indicated by the co-localized ^19^F/^12^C and ^13^C/^12^C signals (Fig. 4B). The ^13^C enrichment in the Thiosymbion regions was similar to that observed in single cell nanoSIMS analyses of the Thiosymbion cells from worm homogenates (Fig. 3A and 3B).

**Fig. 4:**
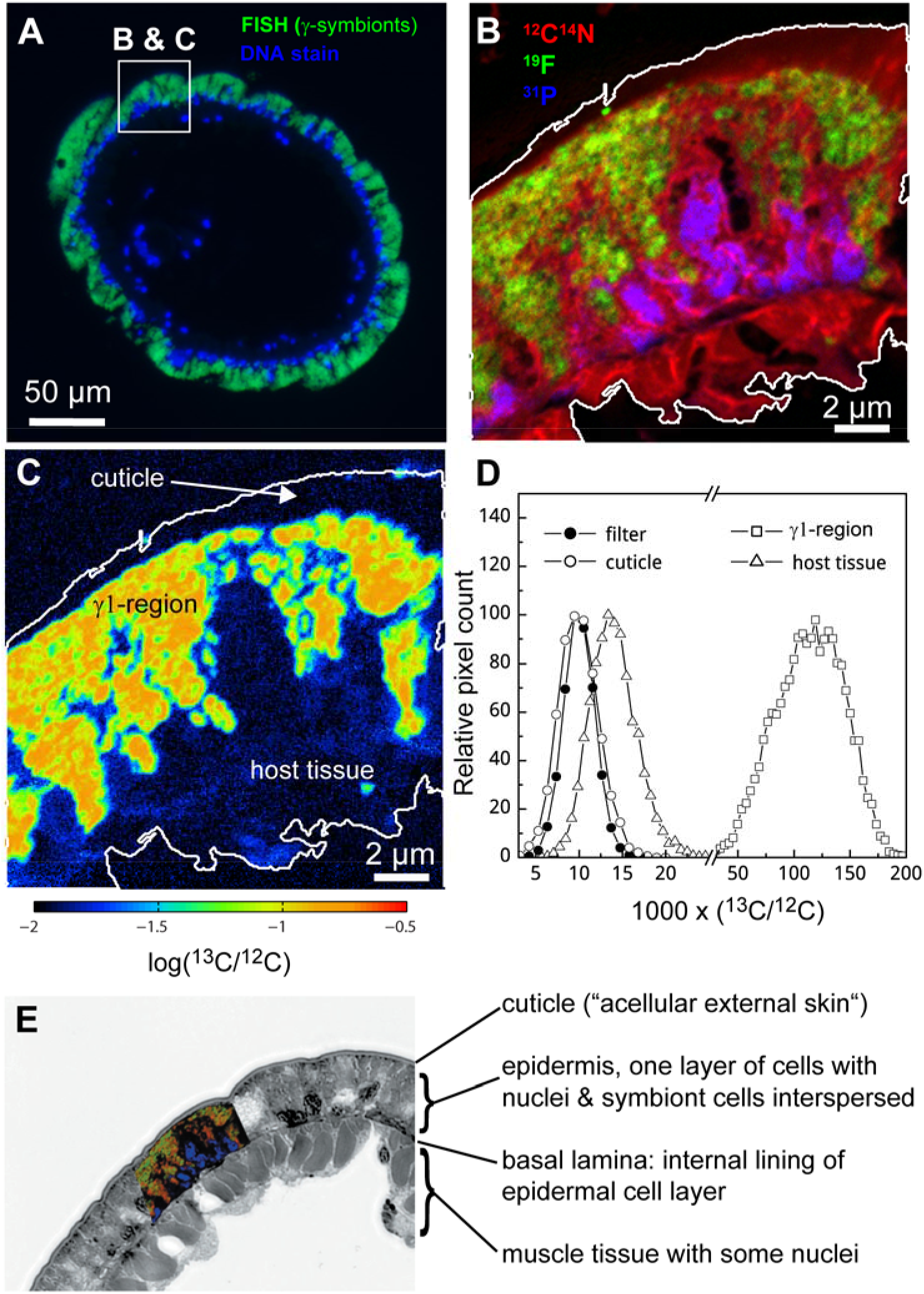
^13^C-enrichment in *O. algarvensis* host tissue and γ1-symbiont (*Ca.* Thiosymbion) region after 25 hour incubation with ^13^C-labelled sodium bicarbonate. A) Epifluorescence image of a cross section of *O. algarvensis* stained with DAPI to visualize DNA and the gammaproteobacteria specific probe GAM42a. B) Composite image of three nanoSIMS images of part of the *O. algarvensis* cross section. The ^31^P channel indicates location of DNA/nuclei. The ^19^F channel is the signal of CARD-HISH hybridization with the gammaproteobacteria specific probe GAM42a. The ^12^C^14^N channel is a general marker of the presence of biomass contrasting the worm section against the carbon only filter onto which the section was mounted. C) NanoSIMS image of the same region as in B) showing the ^13^C/^12^C ratio image. Different regions of the image are labeled based on the information from image B). D) Histograms of the ^13^C/^12^C ratio in these regions. E) Composition of the nanoSIMS and a transmission electron microscope images, putting the nanoSIMS image into the context of ultrastructural features of *O. algarvensis*.

^13^C enrichment was also observed in regions without ^19^F signal where no Thiosymbion cells were present (Fig. 4 C&D). Some enrichment was already observed after 2 hours in these host tissue regions (Fig. 3b). After 25 hours of incubation, the ^13^C abundance in host tissue regions reached 0.0129 (SE=0.0001, N=7 sections from 2 worm individuals; Fig. 3c), and was significantly enriched (ANOVA, p=4×10^-5^).

### *S^0^ content of whole worm and* Ca. *Thiosymbion cells*

Sulfur content varied greatly between individual worms (Table S2). At the beginning of the incubation the average sulfur content was 212.7 nmol per average worm (SD 96.5, N=20) and at the end of the incubation after 25 hours it had decreased by nearly 90% to 28.7 nmol (SD 27.6, N=8). Based on the modeled *Ca*. Thiosymbion cell number (3.33*10^7^) and the fact that *Ca*. Thiosymbion is the only symbiont known to store sulfur (Woyke et al., 2006; Kleiner et al., 2012b), the initial average sulfur content of *Ca*. Thiosymbion was 6.39 fmol cell^-1^ and the cellular sulfur oxidation rate was 0.221 fmol cell^-1^ h^-1^. The *Ca*. Thiosymbion sulfur content falls in the same range as that of the sulfur-oxidizing symbionts of the lucinid clam *Codakia orbicularis* which were previously shown to contain between 1.6 to 32 fmol cell^-1^ (Caro et al., 2007)and the free-living SOX *Allochromatium vinosum*, which had a maximum sulfur content of 33 fmol cell^-1^ (Guerrero et al., 1984).

### Approximately 120 million symbiont cells in an average worm account for 40% of the total holobiont carbon

Based on the difference in ^13^C enrichment between whole worms (SI Results, Table S3) and *Ca*. Thiosymbion cells after 25 hours *Ca*. Thiosymbion contained 33% of the total carbon in the symbiosis (1.65 µmol C). Based on an average biovolume of the *Ca*. Thiosymbion population of 0.112 µl (SI Results), the resulting average carbon to volume ratio for the *Ca*. Thiosymbion was 176.8 fg C µm^-3^. Fitting the model with these data allowed estimating the fraction of carbon contained in all symbionts, which was 40% (Table 1 and SI Results), and absolute symbiont numbers contained in an average worm to be 3.33*10^7^ *Ca*. Thiosymbion cells, 2.62*10^7^ γ3 cells, 2.62*10^7^ δ1 cells, 2.62*10^7^ δ4 cells, and 9.5*10^6^ spirochaete cells.

**Table 1:** Modelled carbon and sulfur budgets and cell numbers. The model also allowed us to extrapolate the development of carbon and sulfur budgets beyond the end of the incubation experiment. The gross carbon fixation rate of *Ca*. Thiosymbion between hours 25 and 125 was only 5 amol carbon cell^-1^ hour^-1^. At 125 hours sulfur was predicted to be entirely consumed; only around 10% of the original PHA was left, and the total carbon content of the holobiont had dropped by 5.6% from the maximum at 25 hours. The carbon loss from the symbiosis between 25 and 125 hours amounted to an average loss of 2.9 nmol carbon hour^-1^. The carbon contained in host tissues had increased from 3 µmol at 0 hours to 3.2 µmol at 125 hours (TABLE 1). The *Ca*. Thiosymbion cell number, which increased from 33 mio at 0 hours to 50 mio at 25 hours, dropped to 47 mio at 125 hours.

| Time (hours) | 0 | 2 | 25 | 125 |
| --- | --- | --- | --- | --- |
| No. of *Ca. Thiosymbion* cells per worm | 3.3E+07 | 4.2E+07 | 5.0E+07 | 4.7E+07 |
| Stored sulfur in *Ca. Thiosymbion* (μmol) | 0.208 | 0.182 | 0.013 | 0 |
| **Fitted model: carbon per symbiosis compartment (μmol)** |  |  |  |  |
| *Ca. Thiosymbion* non-PHA | 1.005 | 1.046 | 1.317 | 1.243 |
| *Ca. Thiosymbion* PHA | 0.647 | 0.616 | 0.364 | 0.067 |
| *Ca. Thiosymbion* total | 1.652 | 1.663 | 1.681 | 1.31 |
| γ3 | 0.203 | 0.202 | 0.192 | 0.157 |
| δ1 | 0.055 | 0.054 | 0.052 | 0.043 |
| δ4 | 0.072 | 0.072 | 0.068 | 0.056 |
| Spirochaete | 0.021 | 0.02 | 0.02 | 0.017 |
| All symbionts | 2.001 | 2.011 | 2.013 | 1.582 |
| Host tissue | 3 | 3.005 | 3.059 | 3.204 |
| Whole worm | 5.001 | 5.016 | 5.072 | 4.786 |
| Seawater | 0.958 | 0.944 | 0.908 | 1.081 |
| **Gross cellular carbon (μmol), modelled without symbiont digestion and PHA remobilization** |  |  |  |  |
| *Ca. Thiosymbion* non-PHA | 1.005 | 1.029 | 1.199 | 1.215 |
| **Apparent (net) carbon fixation rates (amol cell<sup>-1</sup> hour<sup>-1</sup>)<sup>c</sup>** |  |  |  |  |
| Overall *Ca. Thiosymbion* carbon fixation rate<sup>a</sup> | 0 | 616 | 375 | 57 |
| Interval *Ca. Thiosymbion* carbon fixation rate<sup>b</sup> | 0 | 616 | 354 | -22 |
| **Actual (gross) carbon fixation rates (amol cell<sup>-1</sup> hour<sup>-1</sup>)<sup>d</sup>** |  |  |  |  |
| Overall *Ca. Thiosymbion* carbon fixation rate<sup>a</sup> | 0 | 360 | 233 | 50 |
| Interval *Ca. Thiosymbion* carbon fixation rate<sup>b</sup> | 0 | 360 | 222 | 5 |

a: For this calculation the initial *Ca. Thiosymbion* cell number of 33.3 mio was used. Changes in cell number were not considered. Rates are calculated as the hourly change between 0 hours and the respective time point.
b: For this calculation the initial *Ca. Thiosymbion* cell number of 33.3 mio was used. Changes in cell number were not considered. Rates are calculated as the hourly change between each time point and the time point before it.
c: The apparent rates are calculated based on net carbon budgets of "*Ca. Thiosymbion* non-PHA" carbon. The apparent rates do not consider that part of the gross carbon fixation by *Ca. Thiosymbion* is removed through cell digestion by the host and dilution of the freshly fixed <sup>13</sup>C with <sup>12</sup>C from remobilized PHA of which in the fitted model 70% are funneled into "cellular" *Ca. Thiosymbion* carbon. Excretion of freshly fixed carbon was set to zero and thus did not influence the budgets.
d: The gross rates are calculated based on "Gross cellular carbon (μmol), modelled without symbiont digestion and PHA remobilization".

### Oxygen respiration rates differ greatly based on worm sulfur content

Oxygen consumption rates of worms with a high amount of stored sulfur (269.2 ± 54.7 nmol S^0^ per average worm) were around nine fold higher than those of worms with little to no stored sulfur (2.9 ± 3.2 nmol S^0^ per average worm) (Table S4). On average high-sulfur worms consumed 445 pmol O_2_ per minute (N=6, SD=132) and low-sulfur worms 50 pmol O_2_ per minute (N=6, SD=19).

### *More than one third of the carbon in* Ca. *Thiosymbion is stored in polyhydroxyalkanoate*

We determined the PHA content of *Ca*. Thiosymbion by purifying the symbionts from worms and extracting their PHA. The average PHA content in the *Ca*. Thiosymbion population of one worm was 0.7 µmol carbon (SD ±0.15 µmol). In other words, 42% of the carbon in the *Ca.* Thiosymbion population was present as PHA, resulting in an average of 14% of the holobiont carbon and. The value of 42% of carbon in PHA for the *Ca.* Thiosymbion is in the range of what has been reported for related, free-living Chromatiaceae bacteria, in which PHA can make up anywhere from 0 to 83% of the cell dry weight (Liebergesell et al., 1991).

### *Estimating the carbon yield stoichiometry (s2c) in* Ca. *Thiosymbion*

We calculated s2c (i.e. mol carbon fixed per mol of sulfur consumed) using the average S^0^ consumption in a worm within 25 hours (184 nmol) and the average ^13^C+^12^C incorporation (185.7 nmol). The average ^13^C+^12^C incorporation was calculated by correcting the average amount of ^13^C incorporation derived from the bulk measurements (130 nmol) with the average fraction of ^13^C in the seawater DIC pool (70%). The resulting fixed CO_2_ to S^0^ ratio was 1.01. This apparent value for s2c can potentially differ from the actual carbon yield stoichiometry of *Ca*. Thiosymbion, because symbiosis internal carbon fluxes that could not be directly included in the calculation may skew the apparent carbon yield stoichiometry (SI Results). To estimate the lowest and highest possible values for s2c, we fitted the model to achieve maximal and minimal values for s2c, while staying within a reasonable range (95% confidence interval) of the averages of experimentally determined values for carbon uptake, sulfur content, and respiration rates (SI Results). The lowest possible value for s2c in *Ca*. Thiosymbion was 0.7 mol CO_2_ per mol S^0^ and the highest 2.2 mol CO_2_ per mol S^0^. The reported s2c values of free-living SOX are much lower, falling in the range of 0.27–0.58 mol CO_2_ per mol sulfur (various reduced sulfur species) for *Thiobacillus* spp. (Kelly, 1999) and 0.08 mol CO_2_ per mol S^0^ for *Acidithiobacillus ferrooxidans* (Ceskova et al., 2002).

### The energy efficiency of carbon fixation driven by sulfur oxidation in Ca. Thiosymbion is very high

To estimate the efficiency with which *Ca*. Thiosymbion uses sulfur derived energy to fix CO_2_ we used the approach of Klatt and Polerecky (2015), which considers the energy producing (sulfur oxidation) and consuming processes (CO_2_ fixation), as well as additional energy costs for the generation of reducing equivalents (NAD(P)H) required for CO_2_ fixation in the CBB cycle (SI Results). NAD(P)H generation in sulfur oxidizers consumes energy because the redox potential of sulfur-derived electrons has to be lowered by reverse electron transport to be able to reduce NAD(P)^+^. We implemented the equations and chemical reactions from Klatt and Polerecky (2015) in R and, for our calculations, we used the CO_2_ to S^0^ ratio of 1.01 measured in our experiments (SI Results, R script available as SI File S1). The resulting overall energy efficiency ‘e’ was 95.3% (Table S5), which is higher than any previously reported value (Klatt and Polerecky, 2015).

### Model predicts that most or all electrons from sulfur are used for O_2_ reduction

Surprisingly, when we fitted the measured *Olavius* oxygen respiration rates with the model, we found that to obtain a good fit of the model to the data we had to assume that almost all the electrons from sulfur in *Ca*. Thiosymbion had to be used for oxygen reduction (Fig. 5, SI Results, Table S1). This was unexpected, because usually in autotrophic SOX a significant proportion of electrons from sulfur are used to reduce CO_2_ for CO_2_ fixation (Kelly, 1999; Klatt and Polerecky, 2015) and are thus not available for oxygen reduction. This indicates that the majority or all of the electrons required for CO_2_ reduction must come from a source other than S^0^ (see Discussion).

**Fig. 5:**
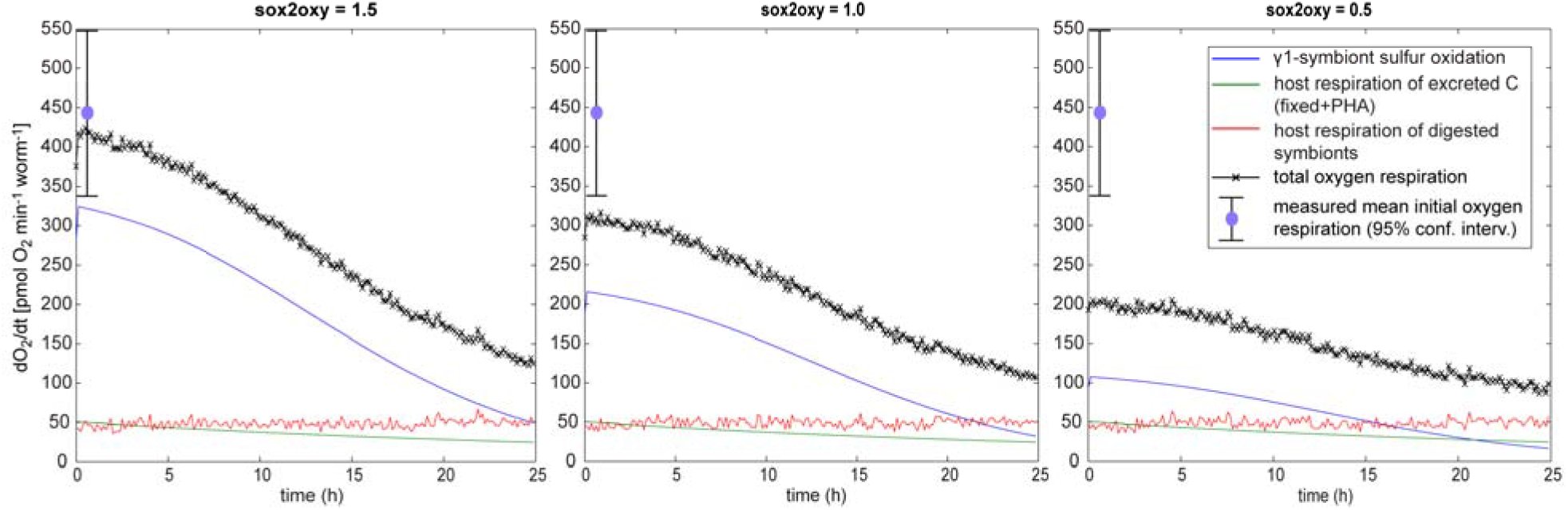
Influence of different respired O_2_ to consumed S^0^ ratios (sox2oxy) on modelled oxygen respiration rates. The mean measured oxygen respiration rate is shown for comparison. A sox2oxy value of 1.5 is the maximum possible value. At sox2oxy = 1.5 all electrons from sulfur are used for oxygen reduction. At sox2oxy values of 1.0 and 0.5, half or all of the CO_2_ fixed by *Ca*. Thiosymbion is reduced with electrons from sulfur thus reducing the amount of electrons used for oxygen reduction.

### Modelled carbon and sulfur budgets in the symbiosis

We used the experimental data to fit the model based on the procedure described in the SI Methods. The obtained fit of the model reflects the experimental data points very well (Fig. 3c) with two exceptions. First, the ^13^C-abundance measurements of single *Ca*. Thiosymbion cells after two hours of incubation could not be fitted with the model. The model predicts a higher ^13^C-enrichment than the one we measured. Part or all of this discrepancy is likely due to isotope loss and dilution of ^13^C in the measured cells due to sample preparation artifacts. Both the loss of freshly synthesized metabolites and the addition of large amounts of ^12^C during the CARD-HISH procedure are known to lead to label loss and dilution (Musat et al., 2014; Woebken et al., 2014; Musat et al., 2016). These label loss and dilution artifacts are expected to be less pronounced at 25 hours due to the overall much higher ^13^C enrichment and conversion of soluble metabolites into “permanent” biomass. Second, the ^13^C-abundance measurements of bulk whole worms after 6, 8 and 16 hours of incubation cannot be satisfactorily fitted with the model. In this case the model predicts slightly higher ^13^C-enrichments than the measured ones (Fig. 3c).

We used the fitted model to determine the per cell carbon fixation rates of *Ca*. Thiosymbion, the sulfur content and the carbon budgets at different time points (TABLE 1). We estimated the carbon fixation rates based on the difference between carbon contents of the *Ca*. Thiosymbion population at given time points excluding the carbon stored in PHA. The maximum net rate was 616 amol carbon cell^-1^ hour^-1^ and the maximum gross rate was 360 amol carbon cell^-1^ hour^-1^ both at the beginning of the experiment. The net rate is higher than the gross rate due to the fact that the model accounts for large amounts of PHA being converted into permanent cellular biomass leading to larger increases in cellular carbon than just by carbon fixation. For the calculation of the gross rate we excluded PHA conversion as well as loss of freshly fixed carbon due to cell digestion by the host. Therefore, the modelled gross rate is likely closer to the true carbon fixation rate. The average gross rate dropped to 222 amol carbon cell^-1^ hour^-1^ between 2 and 25 hours. The doubling time for *Ca*. Thiosymbion population would be 84 hours based on the initial gross rate (SI Discussion). The average carbon gain of the holobiont was 7.5 nmol hour^-1^ in the first two hours and dropped to an average of 2.4 nmol hour^-1^ between 2 and 25 hours.

## Discussion

### High variability of carbon assimilation in individual cells and drop in assimilation rate

There was a high variability in the uptake of inorganic carbon between individual *Ca*. Thiosymbion cells. ^13^C-abundances of individual *Ca*. Thiosymbion cells ranged from 0.0261 to 0.138 in worms from the 25 hour time point (Fig. 3a). While a similar variability in ^13^C-enrichment has been observed in free-living bacteria, we did not expect such variability in the *Olavius* symbionts given the key role that carbon fixation plays in these nutritional symbioses. Metabolic variability between cells from the same bacterial population has been described, for example in bacterial cultures ((Strovas et al., 2007) and references therein), in fresh-water bacteria (Musat et al., 2008), and in sulfur-oxidizing symbionts of a lucinid clam (Caro et al., 2007). One explanation for metabolic variability within a population is genomic heterogeneity (Jaspers and Overmann, 2004). However, the *Ca*. Thiosymbion are obligate endosymbionts and are genetically more homogenous than free-living bacterial populations, as indicated by very low levels of nucleotide polymorphisms in their genomes (Woyke et al., 2006). It is unlikely that diffusive gradients in oxygen (the electron acceptor for the *Ca*. Thiosymbion) from the exterior to the interior region of the worm’s body wall caused the observed metabolic variability. Our nanoSIMS measurements on tissue sections showed that there were no gradients in carbon uptake across the symbiont-containing region (Fig. 4). Alternative explanations for metabolic variability include age and life history (Brookes et al., 2000; Strovas et al., 2007), as well as random fluctuations in the transcription and translation of genes (Taniguchi et al., 2010), all of which could have led to differences in the physiological state of the cells, such as internal sulfur content. Unfortunately, a reliable determination of the sulfur content of individual *Ca*. Thiosymbion cells with the NanoSIMS was not possible as the HISH procedure led to loss of stored sulfur. However, the model allowed us to test for the effect that variability in the amount of stored sulfur in individual cells would have on final ^13^C-enrichment of individual cells. We included a parameter in our model allowing to simulate variable sulfur contents in *Ca*. Thiosymbion cells at the beginning of the experiment. Variability in storage compound content indeed led to a much broader distribution of ^13^C-enrichment values of individual cells (Fig. S1). This suggests that uneven accumulation of stored sulfur in *Ca*. Thiosymbion cells under anoxic conditions could indeed explain part of the uneven ^13^C-enrichment. Interestingly, even when we modelled ^13^C-enrichment in individual cells assuming that all cells start out with the same amount of stored sulfur (Fig. S1), the modelled values still covered a range of ^13^C-enrichments, albeit a narrower range. This is likely due to the inherent assumption of our model that cells are at different stages in their lifecycle at the beginning of oxygen exposure and thus have different initial total carbon contents and divide at different times.

The rate of ^13^C-enrichment in *Ca*. Thiosymbion cells as well as in bulk worms decreased during the incubation experiments (Fig. 3). This was expected given that CO_2_ exchange with the atmosphere was not prevented. However, even though atmospheric exchange was included in the model, the experimental data could not be consistently fitted unless at least one additional parameter leading to a decrease of the holobiont carbon fixation rate down to ∼50% of the initial value was assumed. It is possible that sulfur oxidation rates and thus carbon fixation rates were down-regulated during the incubation, for example because decreasing sulfur concentrations in the *Ca*. Thiosymbion cells caused a down regulation of carbon fixation. Introduction of a parameter in the model that downregulated sulfur oxidation based on decreasing sulfur content led to a good fit of the model with the data. Alternatively, limitation by transport of oxygen through the diffusive boundary layer around the worm could also provide a possible explanation for a drop in rates.

### Why is the energy efficiency of sulfur oxidation based carbon fixation in Ca. Thiosymbion so high?

To our knowledge, the apparent 95% energy efficiency of sulfur oxidation driven CO_2_ fixation in *Ca*. T. algarvensis is the highest ever reported for chemolithoautotrophic SOX. This value is 4 to 24 fold higher than the previously reported values for free-living SOX of 3.91 to 25.3% and 1.2 to 5.3 fold higher than the values reported for symbiotic SOX 18.13 to 77.27% (Table 1 in (Klatt and Polerecky, 2015))(Kelly, 1999). In fact, the efficiency in *Ca.* T. algarvensis is unrealistically high for at least three reasons. First, factorization (i.e. analysis of the partial efficiencies that lead to the overall efficiency ‘e’) of ‘e’ showed that some partial efficiencies of ‘e’ e.g. for sulfur oxidation coupled to terminal electron acceptor reduction (e_SO_) were higher than 100%, which is thermodynamically impossible, as this would imply that energy is generated in this energy requiring process (aka perpetual motion machine) (Table S5, SI Results and Discussion). This suggests that at least one additional factor is missing in the overall chemical reactions used to generate the equations (SI Results and Discussion). The second reason why the efficiency in *Ca*. T. algarvensis is unrealistically high, is that it has been shown that the key enzyme of the CBB cycle, the ribulose bisphosphate carboxylase/oxygenase (RubisCO) is inefficient when oxygen is present, with potentially losses in the range of 21 to 50% of all CO_2_ assimilated (Peterson, 1983; Sharkey, 1988; Cegelski and Schaefer, 2006). Third, overall efficiency of energy transfer to the CBB cycle should be reduced due to energy consuming basic cell maintenance processes and biosynthesis steps, which convert hexoses produced by the CBB cycle into “permanent” biomass such as cell wall components, DNA and proteins.

So how can we explain the extremely high apparent energy efficiency in *Ca.* Thiosymbion in the light of these arguments? There are two potential explanations for this that are not mutually exclusive. The first is that there may be factors leading to higher energy efficiency in *Ca.* Thiosymbion. Three such factors were previously discussed in the literature for symbiotic sulfur-oxidizing bacteria (SOX). First, it was recently suggested that *Ca.* Thiosymbion and most other sulfur-oxidizing symbionts use a modified version of the CBB cycle that is more energy efficient (Kleiner et al., 2012a; Kleiner et al., 2012b). The presence of this modified pathway would, however, not explain the difference in efficiency between free-living and symbiotic sulfur oxidizers, because at least one of the free-living sulfur oxidizers for which energy conversion efficiencies have been reported, *Thiobacillus denitrificans*, was also predicted to use this modified CBB cycle (Kelly, 1999; Kleiner et al., 2012b; Klatt and Polerecky, 2015). *T. denitrificans* has one of the highest reported energy conversion efficiencies among free-living sulfur oxidizers (e = 17.59%, aerobic thiosulfate oxidation) (Klatt and Polerecky, 2015), which is consistent with the use of a more energy-efficient CBB cycle. Second, Klatt and Polerecky (2015) speculated that high efficiency is selected for in symbiotic SOX as compared to free-living SOX, because they likely have evolved highly efficient electron acceptor use due to competition with the host for O_2_ and the fact that they do not have to retain metabolic flexibility in the presumably stable host environment (Klatt and Polerecky, 2015). Third, the host-internal environment inhabited by *Ca.* Thiosymbion may reduce the known inefficiency of RubisCO by reducing oxygen concentrations and providing higher CO_2_ concentrations. It has been shown for several other bacteria-animal symbioses with symbiotic SOX that the host expresses abundant carbonic anhydrases to increase CO_2_ concentrations and facilitate CO_2_ transport to their symbionts (De Cian et al., 2003; Yellowlees et al., 2008; Childress and Girguis, 2011; Hongo et al., 2013; Wippler et al., 2016; Ponnudurai et al., 2017).

The second explanation for the high apparent energy efficiency is that there is a parameter that we did not consider in our calculations. Efficiency calculations so far have been based on the assumption that all energy and reducing equivalents are derived from sulfur oxidation. However, if an additional source of ATP or reducing equivalents existed efficiencies would appear to be higher than they actually are. Currently, we do not have any indications for an additional ATP source in *Ca*. Thiosymbion under the incubation conditions used. However, our result that most or all electrons from sulfur are used for oxygen reduction and are thus not available for CO_2_ reduction (Fig. 5), strongly suggests that an additional source of electrons must contribute to CO_2_ fixation. We found such a potential extra source of reducing equivalents in this study, which is the large amount of PHA stored in *Ca*. Thiosymbion. PHA has been predicted to be produced by *Ca*. Thiosymbion under anaerobic conditions from host fermentative waste (Kleiner et al., 2012b) and is likely remobilized under aerobic conditions (Martin et al., 2006). When PHA is remobilized to funnel carbon back into central carbon metabolism reducing power in form of NAD(P)H is released (Martin et al., 2006). Use of PHA remobilization-derived NAD(P)H for CO_2_ fixation would alleviate the need for reverse electron flow and thus save sulfur-derived energy during CO_2_ fixation. Additionally, electrons from sulfur would not be needed anymore to reduce NAD(P)^+^ and could thus also be used for ATP generation.

To test how partial or complete provision of electrons for CO_2_ reduction from PHA would change the estimated energy efficiency of sulfur oxidation-based carbon fixation in *Ca*. Thiosymbion, we modified the equations of Klatt and Polerecky (2015) to allow for reducing equivalents to be provided from other sources than S^0^ (SI R script File S1, SI results, Table S5). For this we added additional parameters to the equations to account for the fraction of S^0^ used as source of reducing equivalents (y) and the amount of reducing equivalents provided from other sources (x). Based on these equations we calculated different scenarios of reducing equivalent supply (Table S5). If we assume that all NAD(P)H for CO_2_ fixation is derived from PHA then the overall efficiency ‘e’ of sulfur-derived energy use for CO_2_ fixation is 11.38% with partial efficiencies for sulfur oxidation and energy transfer greater than 20% i.e. energy loss or use for cell maintenance in each of these two steps is <80%. If we assume that half of the CO_2_ is reduced with NAD(P)H derived from PHA then the overall efficiency ‘e’ is 32% with a partial efficiency for sulfur oxidation greater than 45%. These efficiencies are still high, but in a more realistic range.

In conclusion, to our knowledge, *Ca. Thiosymbion* has the highest reported energy conversion efficiency for CO_2_ fixation using the existing framework for calculating the efficiency of sulfur-driven carbon fixation (Klatt and Polerecky, 2015). However, using our data, model and additional calculations we show that in *Ca. Thiosymbion* some of the assumptions of the Klatt and Polerecky framework do not hold, because the presence of a sulfur-independent source of reducing equivalents namely PHA boosts the apparent sulfur-derived energy conversion efficiencies. It is likely that for other SOX especially symbiotic SOX a similar use of an additional reducing equivalent source has led to overestimation of energy efficiencies. One factor that we can currently not account for, is the amount of energy that *Ca.* Thiosymbion invests under anoxic conditions to produce the PHA from host-derived waste product and potentially sulfide derived electrons. PHA under oxic conditions likely “subsidizes” sulfur oxidation-driven carbon fixation and therefore the energetic cost for PHA production would need to be included in future efficiency estimates.

### Carbon transfer, heterotrophic carbon fixation and fate of storage compounds

One of the main physiological questions in chemosynthetic symbioses is how carbon is transferred from the symbionts to the host. Two mechanisms have been suggested for transfer of carbon in chemosynthetic symbioses (Distel and Felbeck, 1988; Cavanaugh et al., 2013), which are translocation of metabolites, also referred to as “milking” of the symbionts, and digestion of symbionts in host cells. A recent study by Wippler et al. (2016) provided strong evidence that the main mode of nutrient transfer in *O. algarvensis* from the symbiont to the host occurs via symbiont digestion.

We used our model to test if the ^13^C-enrichment in host tissue that we observed with the nanoSIMS after 2 and 25 hours of incubation could be attributed to transfer of carbon from the symbionts to the host. For this we implemented both milking and digestion as pathways of carbon transfer in our model. One additional parameter implemented in the model that turned out to be critical was heterotrophic carbon fixation by the host tissue. This kind of carbon fixation is due to carboxylation reactions, which are part of intermediary metabolism in most organisms (Wood et al., 1941; Wood et al., 1945). It has been previously shown that in chemosynthetic animals, as in all other animals, significant amounts of inorganic carbon are fixed by the host tissue (Distel and Felbeck, 1988) and for bacteria it has been shown that 3 to 8% of their organic carbon may originate from carboxylation reactions (Feisthauer et al., 2008). We implemented heterotrophic carbon fixation by the host tissue as a percentage of host respiration, as at least for bacteria it has been shown that heterotrophic carbon fixation correlates with respiration. In bacteria heterotrophic carbon fixation amounts to 1-5% carbon fixed per carbon respired (Miltner et al., 2005). As heterotrophic fixation is a direct result of growth, the fixation/respiration ratio is likely much lower in animals, where there is more respiration and less growth.

Using our model we found that digestion and heterotrophic fixation individually, as well as combinations of the two could explain the ^13^C-enrichment. However, to explain ^13^C-enrichment in host tissue solely by heterotrophic fixation, we had to make some extreme assumptions including that the heterotrophic fixation/respiration ratio was very high (0.11) and that part of the PHA remobilized by *Ca*. Thiosymbion would be directly transferred to the host by excretion. In a more realistic scenario with a heterotrophic fixation/respiration ratio of 0.04, 96% of the observed ^13^C-enrichment after 25 hours was due to heterotrophic fixation and 4% due to transfer of freshly fixed carbon from the symbionts. The amount of ^13^C-enrichment in the host tissue contributed by the symbionts may appear small, however, it has to be kept in mind that even after 25 hours only around 9% of the symbiont carbon consist of ^13^C and up to 25 hours it was less than 9%. This means that the bulk of the carbon transferred from the symbionts to the host does not appear in the ^13^C-enrichment measurements. In summary, the largest share of the experimentally observed ^13^C-enrichment in host tissue after 2 and 25 hours of incubation is likely due to transfer of freshly fixed carbon from the symbionts to the host by symbiont digestion. Heterotrophic fixation without carbon transfer from the symbionts is a very unlikely explanation for the ^13^C-enrichment in host tissue because influx of organic carbon is needed to fuel metabolism and respiration in the host tissue to allow for heterotrophic fixation. Future, pulse-chase experiments with ^13^C-bicarbonate will allow to quantify the amount and timescale of carbon transfer from the symbionts to the host more accurately.

### Life style predictions based on the experimental and modelled data

Based on the experimental and modelled data we can make some predictions about the lifestyle of the *O. algarvensis* symbiosis. First, we predict that to achieve maximal growth rates the worms must spend only short periods of time in the oxic layers of the sediment and return quickly to the anoxic layers to enable replenishing of storage compounds in *Ca*. Thiosymbion. This prediction is based on the relatively quick drop (within hours) of the carbon fixation rate of *Ca*. Thiosymbion after exposure of the worms to oxic conditions, which is likely due to depletion of stored sulfur in individual cells. This prediction is supported by the fact that a large majority of worms are usually found in the anoxic layers of the sediment (C. Lott, unpublished).

Second, we can make some predictions on how quickly a worm would deplete its nutritional resources by symbiont digestion if it remained in the oxic layers of the sediment for extended periods of time. Such an extended stay in the oxic layers may occur during the reproductive season in the summer. In the oxic layers of the sediment access to external energy sources for the symbionts would be limited or absent. We used the model to extrapolate the carbon content of the complete symbiont population in one worm over longer periods of oxic conditions (Fig. S2). The model predicted that after 3 weeks roughly half of the symbiont carbon has been consumed and after 6 weeks only around 12% of the original symbiont population carbon remains. This suggests that the worms can persist for several weeks without supply of external energy sources, which matches our observations from long-term storage experiments.

### Experimental and computational methods

See supporting information.

## Supporting information

Matlab Model and Initial Conditions

Suppl. File S1 R-script

## Acknowledgements

We thank the team of the HYDRA Institute on Elba for their support with sample collection and onsite experiments; Jörg Ott, Charles Fisher, Judith Klatt, Gaute Lavik, Florin Musat, Friedrich Widdel and Marc Strous for fruitful discussions on experimental questions, carbon flow and bioenergetics in chemosynthetic symbioses; Tomas Vagner for assistance with NanoSIMS measurements; Sten Littmann for evaluation of NanoSIMS data; Thomas Max, Mike Formolo, and Duygu S. Sevilgen for technical assistance. Funding for this study was provided by the Gordon and Betty Moore Foundation through Grant No. GBMF3811 to ND, the Max Planck Society (ND), the USDA National Institute of Food and Agriculture Hatch project 1014212 (MK), and the U.S. National Science Foundation (grants OIA 1934844 and IOS 2003107 to MK).

