## Supplementary material for "Mechanism of high energy efficiency of carbon fixation by sulfur-oxidizing symbionts revealed by single-cell analyses and metabolic modeling": Matlab Model and Initial Conditions: cell_count.m

```
function [Ng Cg]=cell_count(g)
c12=g.c12; c13=g.c13; phb12=g.phb12; phb13=g.phb13;
Ng=zeros(1,size(c12,2));
Nt=size(c12,2);
for k=1:Nt
    ind=find((c12(:,k)+c13(:,k))>0);
    Ng(k)=length(ind);
end;
Cg = sum(c12+c13+phb12+phb13,1);
```
