## Supplementary material for "Mechanism of high energy efficiency of carbon fixation by sulfur-oxidizing symbionts revealed by single-cell analyses and metabolic modeling": Matlab Model and Initial Conditions: deal2mat.m

```
function y=deal2mat(x,fld)
N=size(x,2);
s=['[y{1:N}]=deal(x.',fld,');'];
eval(s);
% if the cells were dividing, the size of the different y's will increase,
% so fill those that are lower than the maximum size with zeros
Nmax = length(y{1});
for k=2:N
    Nmax = max([Nmax length(y{k})]);
end;

for k=1:N
    Nk=length(y{k});
    if Nk<Nmax
        y{k}=[y{k}; zeros(Nmax-Nk,1)];
    end;
end;
y=cell2mat(y);
```
