## Supplementary material for "Mechanism of high energy efficiency of carbon fixation by sulfur-oxidizing symbionts revealed by single-cell analyses and metabolic modeling": Matlab Model and Initial Conditions: exp_data_olavius.m

```
function exp_data_olavius(f)

ms=12;

my_figure(f);

% gamma1 cells
g1=[0	0	0.011	2E-4
0.1667	10	0.0124	7E-4
2	120	0.01835	0.004
25	1500	0.08982	0.02009];
subplot(2,2,1); hold on;
plot(g1(:,1),g1(:,3),'ro','MarkerSize',ms);

% whole-worm
ww=[0	0	0.0111
0	0	0.0112
0	0	0.0109
0.16667	10	0.01201
0.16667	10	0.0117
0.16667	10	0.01228
0.16667	10	0.01152
0.16667	10	0.01179
0.33333	20	0.01193
0.33333	20	0.01214
0.33333	20	0.01206
0.33333	20	0.0121
0.33333	20	0.01177
0.5	30	0.01241
0.5	30	0.01165
0.5	30	0.0121
0.5	30	0.01245
0.5	30	0.01251
1	60	0.0134
1	60	0.01409
1	60	0.01333
1	60	0.01352
1	60	0.01388
2	120	0.01728
2	120	0.01526
2	120	0.01493
2	120	0.01715
2	120	0.015
4	240	0.02053
4	240	0.02049
4	240	0.0185
4	240	0.02023
4	240	0.01724
6	360	0.01969
6	360	0.01859
6	360	0.02304
6	360	0.02058
6	360	0.02033
8	480	0.02367
8	480	0.0216
8	480	0.02046
8	480	0.01832
8	480	0.02251
16	960	0.02427
16	960	0.02303
16	960	0.03113
16	960	0.02838
25	1500	0.03852
25	1500	0.03331
25	1500	0.04398
25	1500	0.0354
25	1500	0.03205];
plot(ww(:,1),ww(:,3),'bx','MarkerSize',ms);

% host tissue
ht=[25	0.0138	0.01361
25.1	0.0132	0.01303
24.9	0.0127	0.01254
25.2	0.0128	0.01264
24.8	0.0131	0.01293
25.3	0.0128	0.01264
24.7	0.013	0.01283];
plot(ht(:,1),ht(:,3),'gs','MarkerSize',ms);

% seawater
sw=[0	0	0.87083
0	0	0.89065
0.16667	10	0.89251
0.16667	10	0.88621
2	120	0.82334
2	120	0.82544
25	1500	0.51233
25	1500	0.51048];
subplot(2,2,2); hold on;
plot(sw(:,1),sw(:,3),'ks','MarkerSize',ms);

subplot(2,2,3); hold on;
plot(g1(:,1),g1(:,3),'ro','MarkerSize',ms);
plot(ww(:,1),ww(:,3),'bx','MarkerSize',ms);

subplot(2,2,4); hold on;
plot(sw(:,1),sw(:,3),'ks','MarkerSize',ms);
```
