## Supplementary material for "Mechanism of high energy efficiency of carbon fixation by sulfur-oxidizing symbionts revealed by single-cell analyses and metabolic modeling": Matlab Model and Initial Conditions: generate_values.m

```
function x=generate_values(m,s,N,distr,cond)
% generate N numbers 
% if distr=='n', generate numbers with poisson distribution with mean
% derived from the mean M and std S of the corresponding normal
% distribution. here, only numbers that additionally satisfy condition cond
% are returned.
% if distr=='u', generate numbers with uniform distribution between m-s and
% m+s 

if distr=='n'
    if m>0
        %x = normrnd(m,s,N);
        % calculate renormalization factor
        a=m/s^2;
        x = random('poiss',a*m,N)/a;
        [n,xout] = hist(x,50);
        nu = n(find(n>0));
        xu = xout(find(n>0));
        nu = nu/sum(nu);
        xu2 = linspace(min(xu),max(xu),50);
        nu2 = spline(xu,nu,xu2);
        nu2 = nu2/sum(nu2);
        F=cumsum(nu2);
        u = rand(N)*(max(F)-min(F)) + min(F);
        y=zeros(N);
        for ii=1:prod(N)
            y(ii) = invF(u(ii),F,xu2);
        end;
        x = y;
        if nargin>4
            % correct values so that condition cond will be satisfied
            ind = find(~eval(cond));
            if ~isempty(ind)               
                ind1=find(eval(cond));
                % choose random elements from the those that do satisfy the
                % condition cond
                xind = round(random('unif',1,length(ind1),length(ind),1));
                x(ind) = x(ind1(xind));
            end;
        end;
    else
        x = zeros(N(1),N(2));
    end;
elseif distr=='u'
    x = random('unif',m-s,m+s,N);
end;
```
