## Supplementary material for "Mechanism of high energy efficiency of carbon fixation by sulfur-oxidizing symbionts revealed by single-cell analyses and metabolic modeling": Matlab Model and Initial Conditions: invF.m

```
function x=invF(u,F,xu)
df = u-F;
i1 = max(find(df>0));
i2 = min(find(df<0));
if isempty(i1)
    x = min(xu);
elseif isempty(i2)
    x = max(xu);
else
    x1 = xu(i1);
    x2 = xu(i2);
    y1 = F(i1);
    y2 = F(i2);
    x = (x2-x1)/(y2-y1)*(u-y1) + x1;
end;
```
