## Supplementary material for "Mechanism of high energy efficiency of carbon fixation by sulfur-oxidizing symbionts revealed by single-cell analyses and metabolic modeling": Matlab Model and Initial Conditions: transform_for_plotting.m

```
function [g1n g3n d1n d4n spirn hn swn symbn]=transform_for_plotting(g1,g3,d1,d4,spir,h,sw)

g1n=transform_cells(g1);
g3n=transform_cells(g3);
d1n=transform_cells(d1);
d4n=transform_cells(d4);
spirn=transform_cells(spir);

hn.c12=deal2mat(h,'c12');
hn.c13=deal2mat(h,'c13');
hn.a = hn.c13./(hn.c13+hn.c12);

swn.c12=deal2mat(sw,'c12');
swn.c13=deal2mat(sw,'c13');
swn.a = swn.c13./(swn.c13+swn.c12);

symbn.c12 = hn.c12 + sum(g1n.c12,1) + sum(g3n.c12,1) + sum(d1n.c12,1) + sum(d4n.c12,1) + sum(spirn.c12,1);
symbn.c13 = hn.c13 + sum(g1n.c13,1) + sum(g3n.c13,1) + sum(d1n.c13,1) + sum(d4n.c13,1) + sum(spirn.c13,1);
symbn.a = symbn.c13 ./ (symbn.c13+symbn.c12);

function g1=transform_cells(g)
N=size(g,2);
g1.c12=deal2mat(g,'c12');
g1.c13=deal2mat(g,'c13');
g1.phb12=deal2mat(g,'phb12');
g1.phb13=deal2mat(g,'phb13');
a=(g1.c13+g1.phb13)./(g1.c13+g1.phb13+g1.c12+g1.phb12);
ind = find(isnan(a));
a(ind)=zeros(size(ind));
g1.a=a;
g1.s=deal2mat(g,'s');

g1.c12tot=g1.c12+g1.phb12;
g1.c13tot=g1.c13+g1.phb13;
```
