## Supplementary material for "Mechanism of high energy efficiency of carbon fixation by sulfur-oxidizing symbionts revealed by single-cell analyses and metabolic modeling": Matlab Model and Initial Conditions: digest_cells.m

```
function [g h sw o2consumed_symbiont_digestion]=digest_cells(g,h,sw,M,dt)
%% Implement digestion of symbiont cells.

%work only with true cells (with C-content > 0)
c=g.c12+g.c13;
ind=find(c>0);

% momentary number of symbiont cells
N=length(ind);

% number of cells that get digested during dt
Ndig = round(N*h.p_gamma1digestion(M)*dt);

if Ndig>0

    % randomly select Ndig cells from the array of true cells
    itmp = round(N*rand(Ndig,1));
    itmp = setdiff(itmp,0);
    idig = ind(itmp);

    %% digestion of these cells will transfer their C and PHB to the host for
    % respiration and assimilation. 
    % note1: the fraction used for respiration will induce anaplerotic
    % C-fixation as well as O2 consumption! 
    % note2: so far it is not clear what really happens to the PHB and S in
    % the digested cells; therefore it is assumed that PHB is transferred
    % to the host, and utilized as if it was the cellular (fresh,
    % remobilized) carbon, readily digestible by the host
    
    % transferred C to the host
    c12trans = sum(g.c12(idig)) + sum(g.phb12(idig));
    c13trans = sum(g.c13(idig)) + sum(g.phb13(idig));
    
    % utilize this carbon by the host
    [h sw o2consumed_symbiont_digestion]=host_activity(c12trans,c13trans,h,sw);
        
    % set the C+PHB+S content of the digested cells to zero, which will
    % effectively exclude them from the "game" in subsequent "turns"
    g.c12(idig)=zeros(size(idig));
    g.c13(idig)=zeros(size(idig));
    g.phb12(idig)=zeros(size(idig));
    g.phb13(idig)=zeros(size(idig));
    g.s(idig)=zeros(size(idig));

else

    %fprintf(1,'Warning: no cell got digested during dt=%.1e\n',dt);
    %fprintf(1,'M=%d, p=%.1e, N=%d\n', M, h.p_gamma1digestion(M), N);
    o2consumed_symbiont_digestion=0;

end;
```
