## Supplementary material for "Mechanism of high energy efficiency of carbon fixation by sulfur-oxidizing symbionts revealed by single-cell analyses and metabolic modeling": Matlab Model and Initial Conditions: divide_grown_cells.m

```
function g = divide_grown_cells(g,cmax)
%% Find cells which grew above cmax 
% and divide their C, PHB and S contents by two.

c12=g.c12; 
c13=g.c13; 
c=c12+c13;
phb12=g.phb12; 
phb13=g.phb13;
s=g.s; 
r_sox=g.r_sox; 
p_phbremob=g.p_phbremob;
sox2oxy = g.sox2oxy;

% indices of cells that grew over the division threshold
ind=find(c>=cmax);

% first create new (doughter) cells
newc12=c12(ind)/2; 
newc13=c13(ind)/2;
newphb12=phb12(ind)/2; 
newphb13=phb13(ind)/2;
news=s(ind)/2;

% keep their activity rates unchanged
newr=r_sox(ind); 
newp=p_phbremob(ind);

% update the old (mother) cells to new (doughter) cells
c12(ind)=c12(ind)/2; 
c13(ind)=c13(ind)/2;
phb12(ind)=phb12(ind)/2; 
phb13(ind)=phb13(ind)/2;
s(ind)=s(ind)/2;

% now add the new (doughter) cells to the array of all cells
g.c12 = [c12; newc12]; 
g.c13 = [c13; newc13];
g.phb12 = [phb12; newphb12]; 
g.phb13 = [phb13; newphb13];
g.s = [s; news];
g.r_sox = [r_sox; newr]; 
g.p_phbremob = [p_phbremob; newp];
g.sox2oxy = sox2oxy;
```
