## Supplementary material for "Mechanism of high energy efficiency of carbon fixation by sulfur-oxidizing symbionts revealed by single-cell analyses and metabolic modeling": Matlab Model and Initial Conditions: exchange_with_atmosphere.m

```
function sw_new = exchange_with_atmosphere(sw,V,dic,tau,a_natural,dt)

% C concentrations in equilibrium with atmosphere
c12eq = V*dic*(1-a_natural);
c13eq = V*dic*a_natural;

% rate of change depends on the difference between the actual C
% concentration and that in equilibrium 
sw_new.c12 = sw.c12 - (sw.c12-c12eq)/tau*dt;
sw_new.c13 = sw.c13 - (sw.c13-c13eq)/tau*dt;

sw_new.c12 = remove_negative_values(sw_new.c12);
sw_new.c13 = remove_negative_values(sw_new.c13);
```
