## Supplementary material for "Mechanism of high energy efficiency of carbon fixation by sulfur-oxidizing symbionts revealed by single-cell analyses and metabolic modeling": Matlab Model and Initial Conditions: fix_excrete_carbon.m

```
function [g h sw o2consumed]=fix_excrete_carbon(g,h,sw,dt)
% implement change in cells X, olavius host and seawater during a
% time-interval dt, due to 
% - X's cell growth
% - X's phb and sulphur depletion
% - change in sw due to X activity
% - change in host due to C transfer from X

%sw1=sw;

if sum(sw.c12+sw.c13)>0

    % momentary 13C abundance in SW
    a_sw = sw.c13/(sw.c12+sw.c13);
    %a_g = g.c13./(g.c12+g.c13);

    %% gamma1 (X):
    % c-fixation from SW, implementing regulation due to S-content
    [rc,rs,ro2]=gamma1_cfix_rate(g);
    c12fixed = rc * (1-a_sw) * dt;
    c13fixed = rc * a_sw     * dt;
	
	% ro2 is the cellular rate of O2 consumption due to Sox (mol O2 per cell per h), 
	% so sum it up and multiply by dt to calculate O2 consumed during dt by Sox of all symbiont cells
    o2consumed_symbiont = sum(ro2) * dt;

else
    c12fixed = 0;
    c13fixed = 0;
	o2consumed_symbiont = 0;

end;

% excretion of freshly fixed C
c12excr = c12fixed * g.fix2excr;
c13excr = c13fixed * g.fix2excr;

% remobilization of PHB and its excretion to the host
c12phbremob_excr = (1-g.phb2gamma1) * g.p_phbremob .* g.phb12 * dt;
c13phbremob_excr = (1-g.phb2gamma1) * g.p_phbremob .* g.phb13 * dt;

% remobilization of PHB and its utilization for symbiont cell growth
c12phbremob_fixed = g.phb2gamma1 * g.p_phbremob .* g.phb12 * dt;
c13phbremob_fixed = g.phb2gamma1 * g.p_phbremob .* g.phb13 * dt;

% removal of S due to respiration, to make the regulation of C-fixation by
% S content possible (in the next time step)
g.s = g.s - rs*dt;
g.s = remove_negative_values(g.s);

% sum contributions that lead to the growth of the symbiont cells
g.c12 = g.c12 + c12fixed + c12phbremob_fixed - c12excr;
g.c13 = g.c13 + c13fixed + c13phbremob_fixed - c13excr;
g.c12 = remove_negative_values(g.c12);
g.c13 = remove_negative_values(g.c13);

% modify seawater due to C-fixation by the symbiont cells
sw.c12 = sw.c12 - sum(c12fixed);
sw.c13 = sw.c13 - sum(c13fixed);
%sw2=sw;

sw.c12 = remove_negative_values(sw.c12);
sw.c13 = remove_negative_values(sw.c13);
%sw3=sw;

% sum contributions that lead to the removal of phb
g.phb12 = g.phb12 - c12phbremob_excr - c12phbremob_fixed;
g.phb13 = g.phb13 - c13phbremob_excr - c13phbremob_fixed;
g.phb12 = remove_negative_values(g.phb12);
g.phb13 = remove_negative_values(g.phb13);

% sum contributions of C transferred to the host
c12trans = c12excr + c12phbremob_excr;
c13trans = c13excr + c13phbremob_excr;

% modify host's and sw's C-contents due to host's activity
[h sw o2consumed_host]=host_activity(c12trans,c13trans,h,sw);
%sw4=sw;

% o2 consumed by this activity
o2consumed.symbiont = o2consumed_symbiont;
o2consumed.host = o2consumed_host;

%testsw(sw);

%a=0;
```
