## Supplementary material for "Mechanism of high energy efficiency of carbon fixation by sulfur-oxidizing symbionts revealed by single-cell analyses and metabolic modeling": Matlab Model and Initial Conditions: gamma1_cfix_rate.m

```
function [rc,rs,ro2]=gamma1_cfix_rate(g)
% implement regulation of the C-fixation rate by gamma1 by the S content
% calculate also the corresponding O2 consumption rate

rs = g.r_sox .* g.s ./ (g.s + g.r_sox_ks);
rc = g.sox2cfix * rs;
ro2 = g.sox2oxy * rs;
a=0;
```
