## Supplementary material for "Mechanism of high energy efficiency of carbon fixation by sulfur-oxidizing symbionts revealed by single-cell analyses and metabolic modeling": Matlab Model and Initial Conditions: host_activity.m

```
function [h sw o2consumed]=host_activity(c12trans,c13trans,h,sw)
%% Host's activity:
% 1. respiration and assimilation of the transferred carbon
% 2. anaplerotic C-fixation 
% 3. consumption of O2 due to respiration

if sum(sw.c12+sw.c13)>0

    % sw 13C-abundance
    a_sw = sw.c13 ./ (sw.c12+sw.c13);

    % transferred C
    c_trans = c12trans+c13trans;
    % respired C
    c_resp = (1-h.beta)*c_trans;
    
    
    h.c12 = h.c12 ;
    h.c13 = h.c13 ;

    sw.c12 = sw.c12 ;
    sw.c13 = sw.c13 ;

    % assimilate a fraction beta of the transferred C to the host's biomass,
    % fix a fraction alpha of the respired C from the SW to the host's
    % biomass by anaplerotic reaction
    h.c12 = h.c12 + sum(h.beta*c12trans) + sum(c_resp*h.alpha*(1-a_sw));
    h.c13 = h.c13 + sum(h.beta*c13trans) + sum(c_resp*h.alpha*a_sw);

    % the fraction 1-beta of the transferred C will be transferred to the
    % sw as DIC; the anaplerotically fixed C will be removed from the SW
    sw.c12 = sw.c12 + sum((1-h.beta)*c12trans) - sum(c_resp*h.alpha*(1-a_sw));
    sw.c13 = sw.c13 + sum((1-h.beta)*c13trans) - sum(c_resp*h.alpha*a_sw);
    sw.c12 = remove_negative_values(sw.c12);
    sw.c13 = remove_negative_values(sw.c13);

	% O2 consumed from the SW due to host respiration
    o2consumed = sum(c_resp * h.hetero_respir2oxy);

else
    
    % transferred C
    c_trans = c12trans+c13trans;
    % respired C
    c_resp = (1-h.beta)*c_trans;
    
	% O2 consumed from the SW due to host respiration
    o2consumed = sum(c_resp * h.hetero_respir2oxy);

    
    % assimilate a fraction beta of the transferred C to the host's biomass,
    h.c12 = h.c12 + sum(h.beta*c12trans);
    h.c13 = h.c13 + sum(h.beta*c13trans);

    % the fraction 1-beta of the transferred C will be transferred to the
    % sw as DIC
    sw.c12 = sw.c12 + sum((1-h.beta)*c12trans);
    sw.c13 = sw.c13 + sum((1-h.beta)*c13trans);

    if sum(sw.c12+sw.c13)>0
        
        % now C in sw should be >0, so we can calculate its 13C-abundance
        a_sw = sw.c13 ./ (sw.c12+sw.c13);

        % fix a fraction alpha of the respired C from the SW to the host's
        % biomass by anaplerotic reaction
        h.c12 = h.c12 + sum(c_resp*h.alpha*(1-a_sw));
        h.c13 = h.c13 + sum(c_resp*h.alpha*a_sw);

        % the anaplerotically fixed C will be removed from the SW
        sw.c12 = sw.c12 - sum(c_resp*h.alpha*(1-a_sw));
        sw.c13 = sw.c13 - sum(c_resp*h.alpha*a_sw);
        sw.c12 = remove_negative_values(sw.c12);
        sw.c13 = remove_negative_values(sw.c13);
        
    end;

end;
```
