## Supplementary material for "Mechanism of high energy efficiency of carbon fixation by sulfur-oxidizing symbionts revealed by single-cell analyses and metabolic modeling": Matlab Model and Initial Conditions: olavius_algarvensis.m

```
function oa=olavius_algarvensis(c, a, beta, p_gamma1digestion,f_anaplerotic,hetero_respir2oxy)
% define properties of the olavius individual

oa.c12 = (1-a)*c;
oa.c13 = a*c;
oa.beta = beta;
oa.p_gamma1digestion = p_gamma1digestion;
oa.alpha = f_anaplerotic;
oa.hetero_respir2oxy = hetero_respir2oxy;
```
