## Supplementary material for "Mechanism of high energy efficiency of carbon fixation by sulfur-oxidizing symbionts revealed by single-cell analyses and metabolic modeling": Matlab Model and Initial Conditions: symbiont.m

```
function symb=symbiont(c, a, s, phb, r_sox, r_sox_ks, ...
			sox2cfix, fix2excr, phb2gamma1, p_phbremob,sox2oxy)
% define properties of the symbiont cell

symb.c12 = c*(1-a);
symb.c13 = c*a;
symb.phb12 = phb*(1-a);
symb.phb13 = phb*a;
symb.s = s;
symb.r_sox = r_sox;
symb.r_sox_ks = r_sox_ks;
symb.sox2cfix = sox2cfix;
symb.p_phbremob = p_phbremob;
symb.fix2excr = fix2excr;
symb.phb2gamma1 = phb2gamma1;
symb.sox2oxy = sox2oxy;
```
