## Supplementary material for "Mechanism of high energy efficiency of carbon fixation by sulfur-oxidizing symbionts revealed by single-cell analyses and metabolic modeling": Matlab Model and Initial Conditions: initial_conditions.m

```
%% definition of parameters for the C-flow model in Olavius algarvensis

% note: if the values for symbionts are defined as vectors, the
% corresponding order is [gamma1, gamma3, delta1, delta4, spirochaete]; if
% there is only one value, it corresponds to gamma1;

%%%%%%%%%%%%%%%%%
%% symbiont cells
%%%%%%%%%%%%%%%%%

%% fraction of C in the whole symbiosis that is due to (all) symbionts 
%%(0.4 is a value that LP calculated based on the 13C abundance in Gamma1
%%and whole worm + biovolume considerations for all symbionts)
f_symbionts = 1*0.4;

%% desired number of gamma1 cells for which the model will be run
% (should be 10000-20000 or so)
N_g1 = 20000;

%% dimensions (in um)
% cell biovolumes are calculated assuming cylinders with hemispherical ends
c_length = [2.65 1 1.6 1 8.50];
c_width =  [1.40 1 0.6 1 0.25];
c_pcabundance = [1*35 27.5 27.5 27.5 10];

%% carbon content
% Calculated based on C density, Cv (in fg C um^(-3)); Cv can be given
% separately for each type of symbiont; 
%The literature value that Manuel chose is 63 fg C um^-3, which is the mean
%from Fagerbakke et al. 1996
%http://www.int-res.com/articles/ame/10/a010p015.pdf
%The value is consistent with Niculina's PNAS paper in whichthey determined
% (?) Cv to be 4.5 fmol C um^-3 for L. purpurea and 6.4 fmol C um^-3 for C.
% clathratiforme, which is 54-76.8 fg C um^-3 and values in the paper
% http://www.int-res.com/articles/ame/13/a013p113.pdf.
Cv = 1 * [176.8 176.8 63 63 63];

%% PHB content and utilization
% Cellular phb content, expressed as a fraction of the total _initial_
% cellular C-content. That is, if a cell has an initial C-content of
% C_init, it's initial PHB content is C_init*PHBC_ratio, where PHBC_ratio
% is itself a random variable.
%Liebergesell et al. 1991: for Chromatiaceae, depending on substrate 
% 0 to 83% of cellular dry weight is PHA, which corresponds roughly to the 
%fraction of cellular carbon that it can make up (PHA consists of C, O 
%and H). Since TEM images (e.g. Dubilier et al. 2001) show that the 
%symbiont is completely filled with PHA a high value can be assumed.
%So according to the literature the range of values can be 0.0 to 0.83 
%(I would go for 0.4 to 0.6)
PHBC_ratio_mean = 0.42*[1 1 0 0 0];
PHBC_ratio_std = 0.5*PHBC_ratio_mean;
% If PHBC is random, sometimes it can be >1 or <0, which would not
% make sense. The condition >0 is satisfied by the way the random numbers
% are generated, but the condition <1 is not. Specify therefore also the
% maximum PHBC_ratio.
PHBC_ratio_max = 0.83;
% Probability of phb remobilization (per h).
%No Literature, but experience tells me that it is gone after 3 to 5 days, 
%so the parameter should be tuned so that PHA is depleted after 4 days. 
%Giere et al. 1988 observed in I. leuk. that worms were pale within 3 to 5 days.
p_phbremob_mean = 0.023 * [1 0 0 0 0];
p_phbremob_std = 0.5 * p_phbremob_mean;
% Fraction of the remobilized phb that stays in the symbiont cells and is
% used for their growth.
phb2gamma1 =0.7*[1 0 0 0 0];

%% sulfur content of gamma1 cells (mol S per cell)
%Caro et al. 2007: Sulfur-oxidizing Codakia orbicularis symbionts have 
%between 32 fmol per cell (>3 um cells) and 1.6 fmol per cell 
%(<1 um cells) of sulfur 
%Mas and Van Gemerden 1995: Maximum sulfur storage in Chromatiaceae 
%24.5E-15 mol per um-3 (~33 fmol per A. vinosum cell)
%Use 20e-15 as standard literature value
s_c_mean = 6.39e-15;
s_c_std = 0.4 * s_c_mean;
% ks value, determining regulation of Sox by S-content (Sox rate drops to
% 1/2 at S-content = r_sox_ks.)
% No literature data, set to close to zero
r_sox_ks = 0.5*3e-15;

%% rates of Sox (sulfur oxidation) for gamma1 cells (mol S per cell per h)
% Lenk et al. 2011: sulfide oxidation rate by cells in marine sediments 3.7 
% fmol cell-1 day-1, 1.1 fmol cell-1 day-1 -> 
% 1.54167E-16 mol/cell*hour or 4.58333E-17 mol/cell*hour
r_sox_mean =2.0*2.21e-16;
r_sox_std = 0.2 * r_sox_mean;

%% conversion factor between Sox and C-fixation rates in gamma1
%Kelly 1999 (see separate calculation document): 4.76 CO2 can be maximally 
%fixed per S (theoretical maximum: Sulfur oxidation to sulfate using oxygen
%as TEA. Calvin cycle for CO2 fixation), however the real energy conversion
%efficiency of CO2 fixation with sulfur is much lower. According to 
%Kelly 1999, the theoretical maximum CO2 fixation based on DeltaG 
%calculations and the actual CO2 fixation in free-living SOX differ 
%enormously. Energy conversion efficiencies range between 5.6?12.2% for 
%different sulfur compounds and SOX species.
sox2cfix =1.00*0.212*4.76;

%% fraction of freshly fixed C that is excreted by gamma1
%Bright 2000, Cavanaugh 2006, Stewart 2005: In Riftia pachyptial carbon is 
%transferred both by excretion (translocation) and digestion. Bright 2000 
%think it is mostly excretion, but they acknowledge that essential 
%nutrients are most likely transferred by digestion. Distel 1988: In 
%lucinid clams no excretion could be shown. Thus based on the literature 
%this value can be anywhere between 0 to close to 1 (A little freshly 
%fixed carbon has to stay in the symbionts for growth if they grow at all)
fix2excr = 0.0;

%% characteristic cell digestion time (h)
% Time after which a non-growing population of cells would decay to half of
% its momentary abundance due to digestion.
% Note: set to a large value to minimize digestion.
%PhD Thesis of Dennis Fink on Bathy mussels: Almost complete digestion of 
%symbiont population within 10 days (96% lost) AND in a second experiment 
%loss of ~75% of the population within 14 days, after 5 day already ~60% 
%loss. So 5 days = 120 hours seems like a reasonable tau_digest to start 
%out with
tau_digest = 340*[1 1 1 1 1 1];

%%%%%%%
%% worm 
%%%%%%%

%% carbon content of the whole-worm symbiosis (mol C per worm)
% measured in this study: mean = 5e-6
c_worm = 5e-6;

%% number of worms in the Petri dish
N_worms = 36; % 36

%% carbon utilization by the worm
% fraction beta of the carbon _transferred_ to the host from the symbionts 
% that is incorporated into the host biomass; the rest (1-beta) is used for
% repiration to gain energy
% Literature: Edwards 1996, Dash 1977, Mishra 1984, Senapati 1983: The 
%reported values for annelids in the literature range from 15-43% depending
%on lifestyle of the respective worm and lifecycle state e.g. worms that 
%have to move a lot respire more and grow less. The literature actually 
%reports this always as transfer of energy, but what they measure is 
%actually biomass increases and respiration. We fixed the value to 20%.
beta=0.3;

%% anaplerotic C-fixation by the host
% For a given amount of carbon _respired_ by the host, C_resp, an
% equivalent of f_anaplerotic*C_resp is fixed from the seawater to the
% host biomass by anaplerotic reaction.
% For Soil bacteria heterotrophic CO2 fixation is in a range of 1-5% of 
%respiration. As heterotrophic fixation is a result of growth it should be
%much lower in animals, where there is more respiration and less growth. 
%Most likely <1% of respiration.
f_anaplerotic=1*0.04;

%%%%%%%%%%%
%% seawater
%%%%%%%%%%%

%% total volume of sw in the Petri dish (L)
% Because the model will run for one worm, the volume of SW will be later
% corrected for the number of worms in the Petri dish
sw_V_true = 1*15e-3; % 15 ml

%% DIC concentration (mol/L)
% measured in this study: mean = 2.2769 mM
sw_dic = 2.3e-3;

%% air-water exchange lifetime (h)
% DOCUMENTATION???
sw_tau = 1e0*61.8;

%%%%%%%%%%%%
%% model run
%%%%%%%%%%%%

%% number of time steps
Nt=200;

%% total duration (h)
Tmax = 25;

%% number of cells that will be randomly selected for final plotting of individual cells
Nc=60;

%% plotting/display options
plot_graphs = 1;
display_budgets = 1;

%% initial 13C-abundances in the different compartments:
a_g1_init = 1.1e-2;
a_g3_init = 1.1e-2;
a_d1_init = 1.1e-2;
a_d3_init = 1.1e-2;
a_d4_init = 1.1e-2;
a_spir_init = 1.1e-2;
a_oa_init = 1.1e-2;
a_sw_init = 0.88;
a_natural = 1.1e-2;

%%%%%%%%%%%%%%%%%%%%%%%%
%% calculated parameters
%%%%%%%%%%%%%%%%%%%%%%%%

%% biovolumes of symbiont cells
c_biovolume = pi * (c_width.^2)/4 .* (c_length-c_width/3);

%% C-content per symbiont cell (fg C per cell)
% note: based on http://www.int-res.com/articles/ame/13/a013p113.pdf,
% the density of C per cell volume, Cv (in fg C um^(-3)), depends on the
% cellular biovolume, V (in um^3), as Cv = 25.73.*V.^(-0.981)
% Cv = 25.73.*c_biovolume.^(-0.981);
Cc = Cv.*c_biovolume;
% recalculate to mol C per cell (molar weight of C = 12 g/mol)
Cc_molpercell = Cc * 1e-15 / 12;

%% compartment biovolume, expressed as a fraction of total symbiont biovolume
tot_va = sum(c_biovolume.*c_pcabundance);
c_relbiovolume = c_biovolume.*c_pcabundance/tot_va;

%% fractional abundances of symbiont cells
fi = c_pcabundance/sum(c_pcabundance);

%% total number of symbiont cells
Nsymb = f_symbionts*c_worm / sum(Cc_molpercell.*fi);

%% abundace of symbiont cells in the whole worm (cells/worm)
c_abund = fi*Nsymb;

% relative C contributions of the compartments to total C-content of the
% symbionts
%tot_Cca = sum(Cc.*c_pcabundance);
%c_relcc = Cc.*c_pcabundance/tot_Cca;

%% range of cellular C-content in which the symbiont cells grow;
% this means: when the C-content of a cell reaches c_max, the cell divides
% into two halves, i.e., its C, PHB and S contents are halved, while its
% activity remains unchanged
c_c = Cc_molpercell/1.5;
c_max = 2*c_c.*(1-PHBC_ratio_mean);

%% probability of cell digestion (per h)
p_gamma1digestion = log(2)./tau_digest;

%% factor to scale C-contents of the whole-worm symbiosis and seawater such
% that in this scaled-down situation the number of gamma1 cells in the worm
% will be N_g1
%scaling_factor = N_g1/c_abund(1);
scaling_factor = N_g1*Cc_molpercell(1) / (f_symbionts*c_worm - Nsymb*sum(Cc_molpercell(2:end).*fi(2:end)));

%% Carbon content of the host without symbionts (mol C worm^{-1})
% true and scaled down for N_g1 cells
oa_c_true = c_worm*(1-f_symbionts);
oa_c = oa_c_true * scaling_factor;

%% SW volume
% scaled down for N_g1 cells and per 1 worm
sw_V = sw_V_true * scaling_factor/N_worms;

%% time step (in h)
dt=Tmax/(Nt-1);
%% parameters relating Sox and C-respiration to O2 consumption
% molar stoichiometry between Sulphur oxidation rate and O2 consumption
% rate (moles of O2 consumed per mole of S oxidized)
sox2oxy = 1.5;

% molar stoichiometry between host's heterotrophic respiration and O2
% consumption (moles of O2 consumed per mole of C respired)
hetero_respir2oxy = 1;

% index that is considered as the minimum time step
min_ind = 2;
```
