## Supplementary material for "Mechanism of high energy efficiency of carbon fixation by sulfur-oxidizing symbionts revealed by single-cell analyses and metabolic modeling": Matlab Model and Initial Conditions: model14.m

```
%% run model

clear all;
initial_conditions;

%% display values of important parameters
if display_budgets
    fprintf(1,'Scaling factor:\t%.4e\n',scaling_factor);
    fprintf(1,'*** Cellular characteristics (g1 g3 d1 d4 spir):\n');
    fprintf(1,'biovolume (um^3):\t');
    fprintf(1,'%.3f\t',c_biovolume);
    fprintf(1,'\nC-content (fmol C per cell):\t');
    fprintf(1,'%.2f\t',Cc_molpercell*1e15);
    fprintf(1,'\nSymbiont abundancies at t=0 (10^6 cells/worm):\t');
    fprintf(1,'%.1f\t',c_abund/1e6);
    fprintf(1,'\n');
end;

%% initialize gamma1 cells
cid=1;
s=generate_values(s_c_mean,s_c_std,[N_g1,1],'n');
r=generate_values(r_sox_mean,r_sox_std,[N_g1,1],'n');
p=generate_values(p_phbremob_mean(cid),p_phbremob_std(cid),[N_g1,1],'n');
c=generate_values(Cc_molpercell(cid),Cc_molpercell(cid)/3,[N_g1,1],'u');
phb_fraction=generate_values(PHBC_ratio_mean(cid),PHBC_ratio_std(cid),[N_g1,1],'n',['x<' num2str(PHBC_ratio_max)]);
c_cell=c.*(1-phb_fraction);
c_phb=c.*phb_fraction;
g1=symbiont(c_cell,a_g1_init,s,c_phb,r,r_sox_ks,sox2cfix,fix2excr,phb2gamma1(cid),p,sox2oxy);

%% initialize gamma3 cells
cid=2;
N_g3 = round(Nsymb*fi(cid)*scaling_factor);
%N_g3 = round(N_g1/c_relcc(1)*c_relcc(cid));
s=generate_values(0,0,[N_g3,1],'n');
r=generate_values(0,0,[N_g3,1],'n');
p=generate_values(p_phbremob_mean(cid),p_phbremob_std(cid),[N_g3,1],'n');
c=generate_values(Cc_molpercell(cid),Cc_molpercell(cid)/3,[N_g3,1],'u');
phb_fraction=generate_values(PHBC_ratio_mean(cid),PHBC_ratio_std(cid),[N_g3,1],'n',['x<' num2str(PHBC_ratio_max)]);
c_cell=c.*(1-phb_fraction);
c_phb=c.*phb_fraction;
g3=symbiont(c_cell,a_g3_init,s,c_phb,r,r_sox_ks,sox2cfix,fix2excr,phb2gamma1(cid),p,sox2oxy);

%% initialize delta1 cells
cid=3;
N_d1 = round(Nsymb*fi(cid)*scaling_factor);
%N_d1 = round(N_g1/c_relcc(1)*c_relcc(cid));
s=generate_values(0,0,[N_d1,1],'n');
r=generate_values(0,0,[N_d1,1],'n');
p=generate_values(p_phbremob_mean(cid),p_phbremob_std(cid),[N_d1,1],'n');
c=generate_values(Cc_molpercell(cid),Cc_molpercell(cid)/3,[N_d1,1],'u');
phb_fraction=generate_values(PHBC_ratio_mean(cid),PHBC_ratio_std(cid),[N_d1,1],'n',['x<' num2str(PHBC_ratio_max)]);
c_cell=c.*(1-phb_fraction);
c_phb=c.*phb_fraction;
d1=symbiont(c_cell,a_d1_init,s,c_phb,r,r_sox_ks,sox2cfix,fix2excr,phb2gamma1(cid),p,sox2oxy);

%% initialize delta4 cells
cid=4;
N_d4 = round(Nsymb*fi(cid)*scaling_factor);
%N_d4 = round(N_g1/c_relcc(1)*c_relcc(cid));
s=generate_values(0,0,[N_d4,1],'n');
r=generate_values(0,0,[N_d4,1],'n');
p=generate_values(p_phbremob_mean(cid),p_phbremob_std(cid),[N_d4,1],'n');
c=generate_values(Cc_molpercell(cid),Cc_molpercell(cid)/3,[N_d4,1],'u');
phb_fraction=generate_values(PHBC_ratio_mean(cid),PHBC_ratio_std(cid),[N_d4,1],'n',['x<' num2str(PHBC_ratio_max)]);
c_cell=c.*(1-phb_fraction);
c_phb=c.*phb_fraction;
d4=symbiont(c_cell,a_d4_init,s,c_phb,r,r_sox_ks,sox2cfix,fix2excr,phb2gamma1(cid),p,sox2oxy);

%% initialize spirochaete cells
cid=5;
N_spir = round(Nsymb*fi(cid)*scaling_factor);
%N_spir = round(N_g1/c_relcc(1)*c_relcc(cid));
s=generate_values(0,0,[N_spir,1],'n');
r=generate_values(0,0,[N_spir,1],'n');
p=generate_values(p_phbremob_mean(cid),p_phbremob_std(cid),[N_spir,1],'n');
c=generate_values(Cc_molpercell(cid),Cc_molpercell(cid)/3,[N_spir,1],'u');
phb_fraction=generate_values(PHBC_ratio_mean(cid),PHBC_ratio_std(cid),[N_spir,1],'n',['x<' num2str(PHBC_ratio_max)]);
c_cell=c.*(1-phb_fraction);
c_phb=c.*phb_fraction;
spir=symbiont(c_cell,a_spir_init,s,c_phb,r,r_sox_ks,sox2cfix,fix2excr,phb2gamma1(cid),p,sox2oxy);

%% initialize host
h=olavius_algarvensis(oa_c,a_oa_init,beta,p_gamma1digestion,f_anaplerotic,hetero_respir2oxy);

%% initialize seawater
sw=seawater(sw_V,sw_dic,a_sw_init);

%% time steps
t=[0:dt:(Nt-1)*dt];

%% consumed oxygen
o2consumed=[];

%g(1)=g1; h(1)=h1; sw(1)=sw1;

%% loop over time to calculate the dynamics of C-flow
for k=1:(Nt-1)
    
    % seawater-atmosphere exchange
    sw_new = exchange_with_atmosphere(sw(k),sw_V,sw_dic,sw_tau,a_natural,dt);    
    
    % activity of gamma1 cells
    [g1_new h_new sw_new o2consumed_gamma1] = fix_excrete_carbon(g1(k),h(k),sw_new,dt);
    
    % activity of gamma3 cells
    [g3_new h_new sw_new o2consumed_gamma3] = fix_excrete_carbon(g3(k),h_new,sw_new,dt);
    
    % activity of delta1 cells
    [d1_new h_new sw_new o2consumed_delta1] = fix_excrete_carbon(d1(k),h_new,sw_new,dt);
    
    % activity of delta4 cells
    [d4_new h_new sw_new o2consumed_delta4] = fix_excrete_carbon(d4(k),h_new,sw_new,dt);
    
    % activity of spirochaete cells
    [spir_new h_new sw_new o2consumed_spiro] = fix_excrete_carbon(spir(k),h_new,sw_new,dt);
    
    % divide cells
    g1_new = divide_grown_cells(g1_new,c_max(1));
    g3_new = divide_grown_cells(g3_new,c_max(2));
    d1_new = divide_grown_cells(d1_new,c_max(3));
    d4_new = divide_grown_cells(d4_new,c_max(4));
    spir_new = divide_grown_cells(spir_new,c_max(5));
    
    % digest cells
    [g1_new h_new sw_new o2consumed_gamma1_digestion] = digest_cells(g1_new,h_new,sw_new,1,dt);
    [g3_new h_new sw_new o2consumed_gamma3_digestion] = digest_cells(g3_new,h_new,sw_new,2,dt);
    [d1_new h_new sw_new o2consumed_delta1_digestion] = digest_cells(d1_new,h_new,sw_new,3,dt);
    [d4_new h_new sw_new o2consumed_delta4_digestion] = digest_cells(d4_new,h_new,sw_new,4,dt);
    [spir_new h_new sw_new o2consumed_spiro_digestion] = digest_cells(spir_new,h_new,sw_new,5,dt);
    
    % fill variables for this step
    g1(k+1) = g1_new;
    g3(k+1) = g3_new;
    d1(k+1) = d1_new;
    d4(k+1) = d4_new;
    spir(k+1) = spir_new;
    h(k+1) = h_new;
    sw(k+1) = sw_new;
    
    % add contributions to consumed oxygen during dt by ...
    % 1.symbiont's Sox
    o2consumed(1,k) = (o2consumed_gamma1.symbiont + o2consumed_gamma3.symbiont + ...
        o2consumed_delta1.symbiont + o2consumed_delta4.symbiont + o2consumed_spiro.symbiont);
    % 2. respiration of transferred freshly fixed C or remobilized PHB
    o2consumed(2,k) = (o2consumed_gamma1.host+ o2consumed_gamma3.host + ...
        o2consumed_delta1.host+ o2consumed_delta4.host + o2consumed_spiro.host);
    % 3. digestion of symbiont cells
    o2consumed(3,k) = (o2consumed_gamma1_digestion + o2consumed_gamma3_digestion + ...
        o2consumed_delta1_digestion + o2consumed_delta4_digestion + o2consumed_spiro_digestion);
    
end;

[g1n g3n d1n d4n spirn hn swn symn]=transform_for_plotting(g1,g3,d1,d4,spir,h,sw);

if display_budgets    
    %display_C_cells(t,g1n,'g1',plot_graphs,scaling_factor);
    display_C_cells(t,g1n,'g1',0,scaling_factor);
    out=display_cs_budgets(t,g1n,g3n,d1n,d4n,spirn,hn,swn,N_worms,...
        scaling_factor,N_g1,plot_graphs);
end;

if plot_graphs
    plot_c_a(t,g1n,g3n,d1n,d4n,spirn,hn,swn,symn,Nc,scaling_factor);
end;

% display oxygen consumtion results
figure(4);
% o2 consumed per min per worm
subplot(1,1,1);
plot(t(1:end-1),1e12*o2consumed/dt/scaling_factor/60,...
    t(1:end-1),1e12*sum(o2consumed/dt/scaling_factor/60,1),'kx-');
ylabel('dO_2/dt [pmol O_2 min^{-1} worm^{-1}]');
legend('symbiont''s Sox','respiration of excreted C (fixed+PHA)','respiration of digested symbionts','total')
xlabel('time (h)');
title('Oxygen consumption rate by the Olavius symbiosis due to...')
fprintf(1,'*** Oxygen consumption rates [pmol O2/min/worm]\n');
fprintf(1,'\t\t\t\t\tOCR(%.1f)\t\tOCR(%.1f)\n',t(min_ind),t(end));
fprintf(1,'symbiont''s Sox\t\t\t\t%.1f\t\t\t%.1f\n',1e12*o2consumed(1,[min_ind (Nt-1)])/dt/scaling_factor/60);
fprintf(1,'respiration of excreted C\t\t%.1f\t\t\t%.1f\n',1e12*o2consumed(2,[min_ind (Nt-1)])/dt/scaling_factor/60);
fprintf(1,'respiration of digested symbionts\t%.1f\t\t\t%.1f\n',...
    1e12*[mean(o2consumed(3,[min_ind:20])) mean(o2consumed(3,(Nt-1)-[0:19]))]/dt/scaling_factor/60);
fprintf(1,'total\t\t\t\t\t%.1f\t\t\t%.1f\n',1e12*sum(o2consumed(:,[min_ind (Nt-1)]),1)/dt/scaling_factor/60);
fprintf(1,'average OCR during %d h due to Sox: %.1f pmol O2 min^{-1} worm^{-1}\n',Tmax,mean(1e12*o2consumed(1,:)/dt/scaling_factor/60));
fprintf(1,'total S consumed in %d h by Sox: %.3f umol S worm^{-1}\n',Tmax,mean(1e12*o2consumed(1,:)/dt/scaling_factor/60)*60*25*1e-6/sox2oxy);
```
