## Supplementary material for "Mechanism of high energy efficiency of carbon fixation by sulfur-oxidizing symbionts revealed by single-cell analyses and metabolic modeling": Matlab Model and Initial Conditions: display_C_cells.m

```
function display_C_cells(t,g,gstr,flag,sf)

ind0=1;
c12=g.c12(:,ind0); 
c13=g.c13(:,ind0); 
c=c12+c13;
phb12=g.phb12(:,ind0); 
phb13=g.phb13(:,ind0); 
phb=phb12+phb13;
ctot=c+phb;
c=c*1e15; phb=phb*1e15; ctot=ctot*1e15;
nz=find(ctot>0);
fprintf(1,'*** Cellular C budget (fmol C per cell) in %s cells:\n',gstr)
fprintf(1,'t\tCc (mean SD)\tPHB (mean SD)\tC (mean SD)\tN-scaled\tN-true\n');
fprintf(1,'%.1f\t%.2f\t%.2f\t%.2f\t%.2f\t%.2f\t%.2f\t%d\t%.2e\n',...
    t(ind0),mean(c(nz)),std(c(nz)),...
    mean(phb(nz)),std(phb(nz)),...
    mean(ctot(nz)),std(ctot(nz)),length(nz)*[1 1/sf]);
if flag
    my_figure(22);
    subplot(2,3,1); hist(c(nz)); xlabel('C-content (fmol C per cell)');
    subplot(2,3,2); hist(phb(nz)); xlabel('PHB-content (fmol C per cell)'); title(sprintf('time %.f',t(ind0)));
    subplot(2,3,3); hist(ctot(nz)); xlabel('C+PHB (fmol C per cell)');
end;

inde=length(t);
c12=g.c12(:,inde); 
c13=g.c13(:,inde); 
c=c12+c13;
phb12=g.phb12(:,inde); 
phb13=g.phb13(:,inde); 
phb=phb12+phb13;
ctot=c+phb;
c=c*1e15; phb=phb*1e15; ctot=ctot*1e15;
nz=find(ctot>0);
fprintf(1,'%.1f\t%.2f\t%.2f\t%.2f\t%.2f\t%.2f\t%.2f\t%d\t%.2e\n',...
    t(inde),mean(c(nz)),std(c(nz)),...
    mean(phb(nz)),std(phb(nz)),...
    mean(ctot(nz)),std(ctot(nz)),length(nz)*[1 1/sf]);
if flag
    my_figure(22);
    subplot(2,3,4); hist(c(nz)); xlabel('C-content (fmol C per cell)');
    subplot(2,3,5); hist(phb(nz)); xlabel('PHB-content (fmol C per cell)'); title(sprintf('time %.f',t(inde)));
    subplot(2,3,6); hist(ctot(nz)); xlabel('C+PHB (fmol C per cell)');
end;
```
