## Supplementary material for "Mechanism of high energy efficiency of carbon fixation by sulfur-oxidizing symbionts revealed by single-cell analyses and metabolic modeling": Matlab Model and Initial Conditions: display_cs_budgets.m

```
function out=display_cs_budgets(t,g1,g3,d1,d4,spir,h,sw,N_worms,scaling_factor,N_g1,flag)

cc_g1=sum(g1.c12+g1.c13,1);
phb_g1=sum(g1.phb12+g1.phb13,1);
c_g1=cc_g1+phb_g1;
s_g1=sum(g1.s,1);
c_g3=sum(g3.c12+g3.c13+g3.phb12+g3.phb13,1);
c_d1=sum(d1.c12+d1.c13+d1.phb12+d1.phb13,1);
c_d4=sum(d4.c12+d4.c13+d4.phb12+d4.phb13,1);
c_spir=sum(spir.c12+spir.c13+spir.phb12+spir.phb13,1);
c_symbionts=c_g1+c_g3+c_d1+c_d4+c_spir;
c_h=h.c12+h.c13;
c_worm=c_h+c_symbionts;
c_sw=sw.c12+sw.c13;

if flag
    
    my_figure(26);
    fprintf(1,'*** Graph of C+S budgets:\n')
    plot_results(t,s_g1,cc_g1,phb_g1,c_g1,c_g3,c_d1,c_d4,c_spir,c_symbionts,c_h,c_worm,c_sw,1,1,3);
    plot_results(t,s_g1,cc_g1,phb_g1,c_g1,c_g3,c_d1,c_d4,c_spir,c_symbionts,c_h,c_worm,c_sw,1/scaling_factor,2,4);
    
end;

% display budgets in the model scaled down to N_g1 cells
if 0
    fprintf(1,'*** Compartment C budgets (mol; scaled down to Ng1=%d cells)***\n',N_g1);
    out = display_results(t,s_g1,cc_g1,phb_g1,c_g1,c_g3,c_d1,c_d4,c_spir,c_symbionts,...
        c_h,c_worm,c_sw,1,'%.3e');
end;

% display budgets in the whole worm
if 1
    fprintf(1,'*** Compartment C budgets (micro-mol) per whole worm ***\n');
    out = display_results(t,s_g1,cc_g1,phb_g1,c_g1,c_g3,c_d1,c_d4,c_spir,c_symbionts,...
        c_h,c_worm,c_sw,1e6/scaling_factor,'%.3f');
end;

function plot_results(t,s_g1,cc_g1,phb_g1,c_g1,c_g3,c_d1,c_d4,c_spir,c_symbionts,c_h,c_worm,c_sw,sf,s1,s2)
% C and S budgets of the gamma1 symbionts, C budgets in all other symbionts
subplot(2,2,s1);
plot(t,sf*c_g1,'rx-',t,sf*c_g3,'go-',t,sf*c_d1,'bs-',...
    t,sf*c_d4,'mv-',t,sf*c_spir,'c+-',...
    t,sf*c_symbionts,'k-',t,sf*s_g1,'k--');
if sf==1
    fprintf(1,'LEGEND (top):\nred: C_g1; green: C_g3; blue: C_d1; magenta: C_d4; cyan: C_spir; -black: C_symb; --black: S_g1\n');
end;
%legend('C_{g1}','C_{g3}','C_{d1}','C_{d4}','C_{spir}','C_{symb}','S_{g1}','location','eastoutside');
if sf==1
    title('scaled down budgets');
else
    title('budgets per whole worm');
end;
ylabel('C or S (mol)');
xlabel('time (h)')
xlim([min(t) max(t)]);

% C budgets in all symbionts, host, whole symbiosis and sw
subplot(2,2,s2);
plot(t,sf*c_symbionts,'rx-',t,sf*c_h,'go-',t,sf*c_worm,'bs-',t,sf*c_sw,'mv-');
%legend('C_{symbionts}','C_{host}','C_{worm}','C_{sw}','location','eastoutside');
if sf==1
    fprintf(1,'LEGEND (bottom):\nred: symbionts; green: host (without symbionts); blue: whole worm; magenta: seawater\n');
end;
ylabel('C (mol)');
xlabel('time (h)')
xlim([min(t) max(t)]);
    
function out=display_results(t,s_g1,cc_g1,phb_g1,c_g1,c_g3,c_d1,c_d4,c_spir,c_symbionts,c_h,c_worm,c_sw,sf,ff)
out(1,1:3)=display_values(t,sf*s_g1,'S','g1',1,ff);
out(2,1:3)=display_values(t,sf*cc_g1,'Cc','g1',1,ff);
out(3,1:3)=display_values(t,sf*phb_g1,'PHBc','g1',1,ff);
out(4,1:3)=display_values(t,sf*c_g1,'C','g1',1,ff);
out(5,1:3)=display_values(t,sf*c_g3,'C','g3',0,ff);
out(6,1:3)=display_values(t,sf*c_d1,'C','d1',0,ff);
out(7,1:3)=display_values(t,sf*c_d4,'C','d4',0,ff);
out(8,1:3)=display_values(t,sf*c_spir,'C','spir',0,ff);
out(9,1:3)=display_values(t,sf*c_symbionts,'C','symb',0,ff);
out(10,1:3)=display_values(t,sf*c_h,'C','host',0,ff);
out(11,1:3)=display_values(t,sf*c_worm,'C','worm',0,ff);
out(12,1:3)=display_values(t,sf*c_sw,'C','sw',0,ff);
out(13,1:3)=display_values(t,sf*c_worm+sf*c_sw,'C','to air',0,ff);
    
function out=display_values(t,x,s,g,flag,f)
dx=x(end)-x(1);
out=[x(1) x(end) dx];
if flag
    fprintf('\t%s(%.1f)\t%s(%.1f)\td%s\td%s/%s(0)x100\n',s,t(1),s,t(end),s,s,s);
end;
fmt=['%s\t' f '\t' f '\t' f '\t%.1f\n'];
fprintf(fmt,g,x(1),x(end),dx,100*dx/x(1));
%fprintf('%s: %s(%.1f) = %.2e\t%s(%.1f) = %.2e\td%s = %.2e\n',...
%    g,s,t(1),x(1),s,t(end),x(end),s,dx);
```
