## Supplementary material for "Mechanism of high energy efficiency of carbon fixation by sulfur-oxidizing symbionts revealed by single-cell analyses and metabolic modeling": Matlab Model and Initial Conditions: plot_a_symbionts.m

```
function plot_a_symbionts(t,g1,g3,d1,d4,spir,Nc,N1,N2)
my_figure(4);
subplot(N1,N2,1);
plot(t,g1.a(ind,:),'.-');
ylabel('g1: A')
xlim([min(t) max(t)])
subplot(N1,N2,2);
s=size(g3.a,1);
ind=round(rand(Nc,1)*s);
ind=setdiff(ind,0);
plot(t,g3.a(ind,:),'.-');
ylabel('g3: A')
xlim([min(t) max(t)])
subplot(N1,N2,3);
s=size(d1.a,1);
ind=round(rand(Nc,1)*s);
ind=setdiff(ind,0);
plot(t,d1.a(ind,:),'.-');
ylabel('d1: A')
xlim([min(t) max(t)])
subplot(N1,N2,4);
s=size(d4.a,1);
ind=round(rand(Nc,1)*s);
ind=setdiff(ind,0);
plot(t,d4.a(ind,:),'.-');
ylabel('d4: A')
xlim([min(t) max(t)])
subplot(N1,N2,5);
s=size(spir.a,1);
ind=round(rand(Nc,1)*s);
ind=setdiff(ind,0);
plot(t,spir.a(ind,:),'.-');
ylabel('spir: A')
xlim([min(t) max(t)])
```
