## Supplementary material for "Mechanism of high energy efficiency of carbon fixation by sulfur-oxidizing symbionts revealed by single-cell analyses and metabolic modeling": Matlab Model and Initial Conditions: plot_as_gamma1_a_host_sw.m

```
function plot_as_gamma1_a_host_sw(t,g1,h,sw,Nc,N1,N2)
my_figure(3);
s=size(g1.c12,1);
ind=round(rand(Nc,1)*s);
ind=setdiff(ind,0);
xlim([min(t) max(t)])
subplot(N1,N2,1);
plot(t,g1.a(ind,:),'-');
ylabel('g1: A')
xlim([min(t) max(t)])
subplot(N1,N2,2);
plot(t,g1.s(ind,:),'-');
ylabel('g1: S')
xlim([min(t) max(t)])
subplot(N1,N2,3);
plot(t,h.a,'-');
ylabel('h: A')
xlim([min(t) max(t)])
subplot(N1,N2,4);
plot(t,sw.a,'-');
ylabel('sw: A')
xlim([min(t) max(t)])

my_figure(6);
N=size(g1.a,2);
ind=round(linspace(1,N,6));
for i=1:6
    subplot(3,2,i);
    hist(g1.a(:,ind(i)),[0:0.002:0.2])
    xlabel('gamma1 - 13C-abundance');
    v=g1.a(:,ind(i));
    i0=find(v>0);
    qv = std(v(i0))/mean(v(i0));
    title(sprintf('t=%dh: SD/mean=%.3f\n',round(t(ind(i))), qv));
    axis tight
end;
```
