## Supplementary material for "Mechanism of high energy efficiency of carbon fixation by sulfur-oxidizing symbionts revealed by single-cell analyses and metabolic modeling": Matlab Model and Initial Conditions: plot_c_a.m

```
function plot_c_a(t,g1,g3,d1,d4,spir,h,sw,symb,Nc,sf)

N1=3;
N2=3;

%% plot C-content in individual symbiont cells
if 1
    % gamma1
    plot_cells(t,g1,'g1',11,N1,N2,Nc);
end;
if 0
    % gamma3
    plot_cells(t,g3,'g3',12,N1,N2,Nc);
    % delta1
    plot_cells(t,d1,'d1',13,N1,N2,Nc);
    % delta4
    plot_cells(t,d4,'d4',14,N1,N2,Nc);
    % spirochaete
    plot_cells(t,spir,'spir',15,N1,N2,Nc);
end;

%% plot number of cells and total C in (scaled down) symbiont populations
if 1
    plot_nc_symbionts(t,g1,g3,d1,d4,spir,sf);
end;

%% plot C-content in the host (without the symbionts) and in the seawater
if 1
    plot_c_host_sw(t,h,sw,2,3);
end;

%% plot 13-C abundance and S-content in randomly selected gamma1 cells, and
% 13C-abundance in the host tissue and in the seawater
if 1
    plot_as_gamma1_a_host_sw(t,g1,h,sw,Nc,2,2);
end;

%% plot 13-C abundance in randomly selected symbiont cells
if 0
    plot_a_symbionts(t,g1,g3,d1,d4,spir,Nc,2,3);
end;

%% plot modeled and experimentally measured 13C-abundance in gamma1 cells, 
% host tissue, whole-worm symbiosis and seawater as a function of time
if 1
    plot_model_exp(t,g1,g3,d1,d4,spir,h,sw,symb);
end;
```
