## Supplementary material for "Mechanism of high energy efficiency of carbon fixation by sulfur-oxidizing symbionts revealed by single-cell analyses and metabolic modeling": Matlab Model and Initial Conditions: plot_c_host_sw.m

```
function plot_c_host_sw(t,h,sw,N1,N2)
my_figure(2);
subplot(N1,N2,1);
plot(t,h.c12,'-')
ylabel('h: C12')
xlim([min(t) max(t)])
subplot(N1,N2,2);
plot(t,h.c13,'-')
ylabel('h: C13')
xlim([min(t) max(t)])
subplot(N1,N2,3);
plot(t,h.c12+h.c13,'-')
ylabel('h: C12+C13')
xlim([min(t) max(t)])
subplot(N1,N2,4);
plot(t,sw.c12,'-')
ylabel('sw: C12')
xlim([min(t) max(t)])
subplot(N1,N2,5);
plot(t,sw.c13,'-')
ylabel('sw: C13')
xlim([min(t) max(t)])
subplot(N1,N2,6);
plot(t,sw.c12+sw.c13,'-')
ylabel('sw: C12+C13')
xlim([min(t) max(t)])
```
