## Supplementary material for "Mechanism of high energy efficiency of carbon fixation by sulfur-oxidizing symbionts revealed by single-cell analyses and metabolic modeling": Matlab Model and Initial Conditions: plot_cells.m

```
function plot_cells(t,g,gstr,fig,N1,N2,Nc)
s=size(g.c12,1);
ind=round(rand(Nc,1)*s);
ind=setdiff(ind,0);

my_figure(fig);
subplot(N1,N2,1);
plot(t,g.c12(ind,:),'-')
ylabel([gstr ': C12'])
xlim([min(t) max(t)])
subplot(N1,N2,2);
plot(t,g.c13(ind,:),'-')
ylabel([gstr ': C13'])
xlim([min(t) max(t)])
subplot(N1,N2,3);
plot(t,g.c12(ind,:)+g.c13(ind,:),'-')
ylabel([gstr ': C13+C12'])
xlim([min(t) max(t)])
subplot(N1,N2,4);
plot(t,g.phb12(ind,:),'-')
ylabel([gstr ': PHB12'])
xlim([min(t) max(t)])
subplot(N1,N2,5);
plot(t,g.phb13(ind,:),'-')
ylabel([gstr ': PHB13'])
xlim([min(t) max(t)])
subplot(N1,N2,6);
plot(t,g.phb12(ind,:)+g.phb13(ind,:),'-')
ylabel([gstr ': PHB12+PHB13'])
xlim([min(t) max(t)])
subplot(N1,N2,7);
plot(t,g.c12(ind,:)+g.phb12(ind,:),'-')
ylabel([gstr ': C12_{tot}'])
xlim([min(t) max(t)])
subplot(N1,N2,8);
plot(t,g.c13(ind,:)+g.phb13(ind,:),'-')
ylabel([gstr ': C13_{tot}'])
xlim([min(t) max(t)])
subplot(N1,N2,9);
plot(t,g.c12(ind,:)+g.c13(ind,:)+g.phb12(ind,:)+g.phb13(ind,:),'-')
ylabel([gstr ': C_{tot}=C+PHB'])
xlim([min(t) max(t)])
```
