## Supplementary material for "Mechanism of high energy efficiency of carbon fixation by sulfur-oxidizing symbionts revealed by single-cell analyses and metabolic modeling": Matlab Model and Initial Conditions: plot_model_exp.m

```
function plot_model_exp(t,g1,g3,d1,d4,spir,h,sw,sym)
for j=1:length(t)
    [m s]=get_mean_std(g1,j);
    m1(j,1)=m; s1(j,1)=s;
    if 0
        [m2 s2]=get_mean_std(g3,j);
        [m3 s3]=get_mean_std(d1,j);
        [m4 s4]=get_mean_std(d4,j);
        [m5 s5]=get_mean_std(spir,j);
    end;
    if 0
        errorbar(t(j),m2,s2,'gv','MarkerSize',8);
        errorbar(t(j),m3,s3,'bo','MarkerSize',12);
        errorbar(t(j),m4,s4,'mx','MarkerSize',16);
        errorbar(t(j),m5,s5,'cs','MarkerSize',20);
    end;
end;

my_figure(25);

subplot(2,2,1);
hold off
%errorbar(t,m1,s1,'r.-'); 
plot(t,m1,'r.-'); 
hold on;
plot(t,h.a,'gs-')
plot(t,sym.a,'b.-')
xlim([min(t) max(t)])
ylim([0.009 0.2]);
ylabel('13C-abundance');
xlabel('time (h)')
title('red: gamma1;  green: host; blue: g1+h');
set(gca,'yscale','log');

subplot(2,2,2); hold off
plot(t,sw.a,'cv-');
xlim([min(t) max(t)])
ylim([0 1]);
xlabel('time (h)')
title('cyan: seawater');

subplot(2,2,3);
hold off
%errorbar(t,m1,s1,'r.-'); 
plot(t,m1,'r.-'); 
hold on;
plot(t,h.a,'gs-')
plot(t,sym.a,'b.-')
ylabel('13C-abundance');
xlabel('time (h)')
title('red: gamma1;  green: host');

subplot(2,2,4); hold off;
plot(t,sw.a,'cv-');
xlim([0 2])
ylim([0 1]);
xlabel('time (h)')
title('cyan: seawater');

ind=find(t>1.9 & t<2.1);
ind=ind(1);
fprintf(1,'*** Modeled 13C-abundances:\n');
fprintf(1,'t\tgamma1\thost\tworm\tsw\n');
fprintf(1,'%.1f\t%.5f\t%.5f\t%.5f\t%.5f\n',...
    t(ind),m1(ind),h.a(ind),sym.a(ind),sw.a(ind));
ind=length(t);
fprintf(1,'%.1f\t%.5f\t%.5f\t%.5f\t%.5f\n',...
    t(ind),m1(ind),h.a(ind),sym.a(ind),sw.a(ind));

exp_data_olavius(25);

subplot(2,2,3);
xlim([0 7])
ylim([0.01 0.03]);
```
