## Supplementary material for "Mechanism of high energy efficiency of carbon fixation by sulfur-oxidizing symbionts revealed by single-cell analyses and metabolic modeling": Matlab Model and Initial Conditions: plot_nc_symbionts.m

```
function plot_nc_symbionts(t,g1,g3,d1,d4,spir,sf)
[Ng1 Cg1]=cell_count(g1);
[Ng3 Cg3]=cell_count(g3);
[Nd1 Cd1]=cell_count(d1);
[Nd4 Cd4]=cell_count(d4);
[Nspir Cspir]=cell_count(spir);
my_figure(1);
subplot(2,1,1);
plot(t,Ng1,'rx-',t,Ng3,'go-',t,Nd1,'bs-',t,Nd4,'mv-',t,Nspir,'c+-');
ylabel('N')
title('Number of symbiont cells (scaled down model)');
legend('gamma1','gamma3','delta1','delta4','spirochaete',...
    'location','eastoutside');
xlim([min(t) max(t)])
subplot(2,1,2);
plot(t,Cg1/sf,'rx-',t,Cg3/sf,'go-',t,Cd1/sf,'bs-',t,Cd4/sf,'mv-',t,Cspir/sf,'c+-');
ylabel('C (mol)');
title('Total C-content in symbiont compartments (per whole worm)')
xlabel('time (h)')
legend('gamma1','gamma3','delta1','delta4','spirochaete',...
    'location','eastoutside');
xlim([min(t) max(t)])        
```
