## Supplementary material for "Mechanism of high energy efficiency of carbon fixation by sulfur-oxidizing symbionts revealed by single-cell analyses and metabolic modeling": Matlab Model and Initial Conditions: startup.m

```
%% startup script

addpath('auxil','fnc','plot');

fprintf(1,'***************************************************\n');
fprintf(1,'Modeling of C-flow in Olavius algarvensis symbiosis\n');
fprintf(1,'L. Polerecky, 28.8.2013, version 13\n'); 
fprintf(1,'***************************************************\n');
fprintf(1,'Edit initial_conditions.m to set the parameters of the model.\n');
fprintf(1,'Run model13 to run the model.\n');

fprintf(1,'Run save_results(out,''filename'') to save C-budgets data in a text file.\n');
fprintf(1,'Run print_results(N,''filename'') to export figure N as EPS.\n');
```
