## Supplementary material for "Mechanism of high energy efficiency of carbon fixation by sulfur-oxidizing symbionts revealed by single-cell analyses and metabolic modeling": Suppl. File S1 R-script

# Functions implementing the formalism described by Klatt & Polerecky (2015) 
# to quantify efficiency of aerobic elemental sulfur oxidation

# Gibbs free energy of formation at standard biochemical conditions, in kJ/mol
dG0 <- list(sulfate = -744.63, 
			water = -237.18, 
			protons = 0, 
			H2S = -27.87,	  # sulfide
			S2O3 = -522.5,	# thiosulfate
			O2 = 16.4, 		  # O2 dissolved in water
			NO3=-111.3,		  # nitrate
			N2 = 18.2, 		  # di-nitrogen
			Corg = -152.57, # CH2O assumed as fructose (C6H1206), the value is normalized per one C
			CO2 = -386.02, 
			S0 = 0, 		    # zero-valent sulfur
			deltaNAD=60.99, # dGf^0(NADH)-dGf^0(NAD+)
			deltaFd=38.88,  # dGf^0(Fdred)-dGf^0(Fdox)
  		deltaFAD=42.65, # dGf^0(FADH2)-dGf^0(FAD)
			EATP = 41,      # Energy content of ATP
  		EATP_Calvin= 3*41, # dE(CO2 fix) in kJ per mole CO2 (3ATP per CO2)
  		EATP_rTCA = 5/3*41, # dE(CO2 fix) in kJ per mole CO2 (1.67ATP per CO2)
			dG_Calvin=69.67, # dG(CO2 fix) in kJ per mole CO2
			dG_rTCA=64.33)   # dG(CO2 fix) in kJ per mole CO2

# universal gas constant, in kJ/mol/K
R <- 0.00831451

#####################################################################
# Aerobic S0 oxidation coupled to CO2 fixation
#####################################################################

# stoichiometric coefficients and other parameters as a function of x and y

# function for calculating the efficiency from known stoichiometry
# - stoichiometry is given by parameter y
# - carbon fixation assumed to be done via Calvin cycle
# - additional arguments include the actual concentrations (in mol/L) of the reactants involved
#   (sulfate, O2, CO2), temperature (in degC), pH, redox carriers (NAD/NADH)
# - default values of the additional arguments correspond to standard biochemical conditions
#   (reactant concentrations 1M, Temperature 25 degC, pH 7)
aerob_efficiency <- function(gamma=1, delta=1, sulfate=1, O2=1, CO2=1, Temp=25, pH=7, NAD=1, NADH=1){
  x <- delta+gamma-3/2
  y <- 1-2*delta/3
  protons <- 10^(-pH)
  # quotients Q
  # energy generating reaction (O2 reduction)
  Q1 <- sulfate * protons^2 * O2^(-1.5)
  # energy conserving reaction (RET)
  Q3 <- sulfate * protons^5 * NADH^3 * NAD^(-3)
  # Gibbs free energies, in kJ per mole S0
  # energy generating reaction (O2 reduction)
  dGr0_O2red <- -1*dG0$S0 + 1*dG0$sulfate - 1*dG0$water + 2*dG0$protons - 1.5*dG0$O2
  # CO2 fixation (for now Calvin cycle)
  #dGr0_CO2fix <- 1.5*dG0$dG_Calvin # kJ per mole S0
  dGr0_CO2fix <- dG0$dG_Calvin # kJ per mole C
  # reverse electron transport
  dGr0_RET <- -1*dG0$S0 + 1*dG0$sulfate - 4*dG0$water + 3*dG0$deltaNAD + 5*dG0$protons # kJ per mole S0
  # total CO2 reduction
  dGr0_CO2red <- (1.5*y+x)*dGr0_CO2fix + y*dGr0_RET # kJ
  # calculate T-correction
  corr_O2red <-  R*(273.15+Temp)*log(Q1)
  corr_CO2red <- R*(273.15+Temp)*log(Q3)
  #Q <- Q1^y * Q3^(1-y)
  # apply T-correction
  dGr_O2red <- dGr0_O2red + corr_O2red
  dGr_CO2red <- dGr0_CO2red + y*corr_CO2red
  # finally calculate the energy conservation efficiencies
  #eff <- -(1-y)*dGr_CO2red / ( y*dGr_O2red )
  eff <- -1*dGr_CO2red / ( (1-y)*dGr_O2red )
  eff_CO2 <- dG0$dG_Calvin/dG0$EATP_Calvin
  #alpha <- dGr0_CO2fix/dGr_CO2red
  alpha <- (1.5)*dGr0_CO2fix/((1.5)*dGr0_CO2fix + dGr0_RET + corr_CO2red)
  factor_min <- (y*(alpha/eff_CO2 + (1-alpha)) + x/1.5*alpha/eff_CO2) / (y + x/1.5*alpha)
  eff_SO_min <- eff*factor_min  
  eff_RET_min <- y*(1-alpha)/(1/eff*(y+alpha*x/1.5) - alpha/eff_CO2*(y+x/1.5))
  # in addition to efficiencies (eff, eff_CO2, eff_RET_min, eff_SO_min), 
  # the function also returns the Gibbs free energies of the energy gaining
  # (dGr_O2red) and energy conserving (dGr_CO2red) parts of the reaction,
  # as well as the quotient Q and coefficient alpha and factor
  return(list(x=x, y=y, eff=eff, eff_CO2=eff_CO2, eff_SO_min=eff_SO_min, eff_RET_min=eff_RET_min, 
              alpha=alpha,factor_min=factor_min, Q1=Q1, Q3=Q3, 
              dGr_RET=dGr0_RET+corr_CO2red, dGr_CO2red=dGr_CO2red, dGr_O2red=dGr_O2red))
}

# calculate efficiency at std conditions
(e<-aerob_efficiency(gamma=1, delta=1.5))
# type e in the R terminal to display the output values

deltas <- seq(0.5,1.5,length.out = 20)
e<-aerob_efficiency(gamma=1, delta=deltas)
effs <- e$eff
effs_SO_min <- e$eff_SO_min
effs_RET_min <- e$eff_RET_min
xs <- e$x
ys <- e$y
plot(deltas, effs, type="l", col='black', xlab='delta', ylab='efficiency', ylim=c(0,1.5))
lines(deltas, effs_SO_min, col='red')
lines(deltas, effs_RET_min, col='green')
lines(deltas, xs, col='blue')
lines(deltas, ys, col='magenta')
legend('topright', legend=c('e-total','e-min(SO)','e-min(RET)','x','y'), bty="n", 
       col=c('black','red','green','blue','magenta'), pch=c(NA,NA,NA,NA,NA),lty=c(1,1,1,1,1))
