## Supplemental Materials for "Mechanism of high energy efficiency of carbon fixation by sulfur-oxidizing symbionts revealed by single-cell analyses and metabolic modeling"

### SI Materials and Methods

#### Incubation experiment

##### *Sampling*

Sand containing *Olavius algarvensis* specimens was collected off the coast of Sant'Andrea, Island of Elba, Italy, at a water depth of about 6 – 7 m by SCUBA diving on January 2nd 2009 and stored in closed 8 l plastic containers. On the day before each incubation worms were carefully separated from the sediment by decantation with seawater and kept overnight in sealed glass vials (Exetainer, Labco Ltd., England) filled with anoxic seawater and sediment from their collection site.

##### *Incubations*

Artificial seawater (ASW: 0.756 mM KBr, 8.05 mM KCl, 10 mM $CaCl_2$, 27.89 mM $MgCl_2$, 27.6 mM $MgSO_4$, 451 mM NaCl, 4.67 µM $NH_4Cl$, 1.47 µM $KH_2PO_4$) was prepared without the addition of carbon sources and kept anoxic after autoclaving. The incubation medium was prepared from this ASW by aerating it and adding $^{13}C$-labelled sodium bicarbonate (98 atom % $^{13}C$, Sigma Aldrich Inc.) as the sole carbon source to trace aerobic inorganic carbon uptake in the incubations. The incubation medium had the same salinity (39 PSU) and pH (8.2) as the seawater at the worm collection site. The temperature was kept at 15° C throughout the incubation, corresponding to the *in situ* temperature.

Incubations were conducted from January 6th to 9th for 10, 20, 30 min, 1, 2, 4, 6, 8, 16, and 25 hours. First, worms were taken from the glass vials and washed out of the sediment using carbon-free anoxic ASW. At the start of the incubation only white and actively moving worms were placed in plastic petri dishes of 85 mm diameter, with a thin layer of glass beads (ø 0.4 - 0.6 mm; Sartorius, Göttingen, Germany) and 15 ml of the incubation medium containing $^{13}C$-labelled bicarbonate. The initial concentration of dissolved inorganic carbon (DIC) and its isotopic composition was determined by GC-IRMS analysis (see below) as 2.3 mM and 88 atom % $^{13}C$, respectively. No external electron donor was supplied.

The Petri dishes were kept open during the incubation and contained on average 36 (35.55) worms. The exact worm number was documented for each dish (Table S6). Independent Petri dishes were prepared for each time point of the incubation. To control for $CO_2$ exchange with the atmosphere Petri dishes without worms were incubated in parallel. To stop the incubation the Petri dish was put on ice, the incubation solution removed from the dish with a syringe and replaced twice with ice-cold carbon-free ASW. Only active worms, as determined by visual inspection, were used for further analyses. All worms were photographed to determine their biovolume.

For the analysis of bulk carbon content and isotopic composition 15 worms were sampled per time point, and frozen in groups of three in five silver cups at -20°C. For nanoSIMS analyses 6 worms were sampled singly and prepared as described below. The rest of the worms from each time point were frozen singly in 0.5 ml Eppendorff cups at -20°C for sulfur content analysis. Worms for time point 0 min were processed immediately after the removal from the glass vials and never brought into contact with the $^{13}C$-containing incubation medium. At time points 0, 10 min, 2 hours and 25 hours samples of the incubation medium were taken and analysed for total DIC concentration and isotopic composition.

##### Analysis of the incubation medium

6 ml of medium were fixed with 100 µl of a concentrated solution of $HgCl_2$ and stored in a sealed glass vial. For analysis 1 ml of the fixed sample was transferred to a 5 ml Exetainer septum vial (Labco Ltd., England) and acidified with 100 µl of 10% $H_3PO_4$. Then $^{12}C$ and $^{13}C$ were quantified in headspace samples using a gas chromatography – isotope ratio mass spectrometer (GC-IRMS, VG Optima, Manchester, UK) as described previously (Holtappels et al., 2011). Pure $CO_2$ gas with a natural abundance of $^{13}C$ was used as a calibration standard.

##### Bulk worm analyses

The $^{12}C$ and $^{13}C$ content of whole *O. algarvensis* worms was quantified in an automated elemental analyzer (Thermo Flash EA, 1112 Series) coupled to a Delta Plus Advantage mass spectrometer (Thermo Finnigan) using standard methods (Musat et al., 2008). Five replicates per time point were analysed, each containing three worm individuals.

##### In situ *hybridization of worm homogenates and tissue sections*

Three of the six worms taken for nanoSIMS analyses were prepared for whole worm sectioning and the other three as homogenates for single cell analyses. For worm sections, the worms were washed immediately after collection in ice-cold sterile-filtered (0.2 µm) seawater (fSW), fixed in 4% paraformaldehyde in fSW at 4°C for 12 hours, and washed three times in fSW. Subsequently, whole worms were transferred to 70% ethanol and stored at 4°C. For homogenate analyses, individual worms were homogenized in 600 µl ice-cold fSW and then centrifuged at 13000 rpm for 5 min. The seawater was decanted, the pellet was resuspended and fixed in 4% paraformaldehyde in fSW for 12 hours and stored in 70% ethanol at –20°C.

Prior to nanoSIMS analysis, whole fixed worms were embedded in Steedman's wax, cut in 5 µm thick sections, the sections placed on gold-palladium-coated polycarbonate filters (Musat et al., 2008) and de-waxed (Pernthaler and Pernthaler, 2005). Aliquots of the homogenates were pipetted directly onto gold-palladium-coated polycarbonate filters.

Catalyzed reporter deposition (CARD) halogen *in situ* hybridization (HISH) on sections and homogenates was performed as described previously (Musat et al., 2008). We used a tyramide synthesized from the fluorine-containing dye Oregon Green® 488-X (Molecular Probes, Inc.), which allows detection of probe binding both with an epifluorescence microscope based on the fluorescent dye and with the nanoSIMS based on the fluorine. The following oligonucleotide probes were used: EUB I-III (Daims et al., 1999) as a positive control, GAM42a (Manz et al., 1992) as probe specific to Gammaproteobacteria, OalgGam1 as probe specific to *Ca.* Thiosymbion (Ruehland et al., 2008), and NON338 (Wallner et al., 1993) as a negative control. The γ-symbionts were discriminated from the δ-symbionts based on the HISH signal and *Ca*. Thiosymbion from the γ3-symbiont based on size (Fig. 2).

##### NanoSIMS analyses of worm homogenates and tissue sections

$^{13}C$ incorporation into worm tissue and single symbiont cells was analyzed by nanometre-scale secondary ion mass spectrometry (nanoSIMS) using a NanoSIMS 50L (CAMECA, Paris, France).

Images recorded simultaneously included the secondary electron (SE) image for the approximate characterization of the sample topology, $^{12}C^{14}N^{-}$ for the localization of biomass, $^{12}C^{-}$ and $^{13}C^{-}$ for the quantification of carbon uptake and transfer, $^{19}F^{-}$ for the identification of HISH-probed symbiont cells, and $^{31}P^{-}$ for the localization of the host cell nuclei (in worm sections only). All samples were pre-sputtered with a $Cs^{+}$ beam of ~500 pA to remove surface contaminations and to implant $Cs^{+}$ ions into the sample surface. For analysis, the samples were sputtered with a 4.7 pA $Cs^{+}$ primary ion beam focused into a spot of ~150 nm diameter that was scanned over the sample with a counting time of 1 ms per pixel.

NanoSIMS data-sets were analysed using the Look@NanoSIMS program (Polerecky et al., 2012). For single cell analyses, regions of interest were drawn based on $^{12}C^{14}N^{-}$ images. Symbiont cells were classified based on their size and co-localization of the $^{12}C^{14}N^{-}$ and $^{19}F^{-}$ signals. The isotope ratio $r = {}^{13}C/{}^{12}C$ was calculated for each individual cell based on the total $^{13}C^{-}$ and $^{12}C^{-}$ counts accumulated over the cell volume. Subsequently, the $^{13}C$ abundance, defined as $A = {}^{13}C/({}^{13}C+{}^{12}C)$, was calculated as $A = r/(r+1)$. Between 13 and 65 individual cells of all identified symbiont types (*Ca*. Thiosymbion, γ3 and δ) were analyzed for each time point. To follow carbon transfer between *Ca*. Thiosymbion cells and host tissues, seven sections of two worms incubated for 25 hours were additionally analysed. Again, tissue regions inhabited by *Ca*. Thiosymbion cells were identified from co-localization of the $^{12}C^{14}N^{-}$ and $^{19}F^{-}$ signals, whereas host tissue without intact *Ca*. Thiosymbion cells was identified as areas with $^{12}C^{14}N^{-}$ and without $^{19}F^{-}$ signals.

#### *S° content analysis*

Frozen worms where thawed in 1000 µl HPLC-grade methanol 99.9% (Carl Roth, Karlsruhe, Germany), smashed with a glass rod, vortexed for 10 s and extracted on a shaker over night at 4°C. The extract was filtered through a 0.45 µm PTFE membrane Acrodisc CR 4mm syringe filter (Pall Life Sciences, USA) into a 1.5 septum glass vial (Zinsser analytic, Frankfurt, Germany). Elemental sulfur content was measured by HPLC as described in Kamyshny et al. 2009 (Kamyshny et al., 2009). The sulfur content of each worm was normalized to its biovolume. To make the sulfur measurements comparable to the carbon bulk measurements, average normalized sulfur contents per time point were multiplied by the average biovolume of the worms used for carbon bulk analyses (0.532 µl = "average worm").

#### *NMR analyses of PHA content in* Ca. *Thiosymbion*

To determine the initial PHA content of *Ca*. Thiosymbion, worms were collected and treated similarly to the incubation experiment. Worms were removed from the sediment and stored overnight in three sealed glass vials (Exetainer, Labco Ltd., England) containing anoxic artificial seawater, a glass bead mix to simulate the sediment and 100 worms each. The next day worms were removed from the vials and homogenized in a Duall® homogenizer (Tissue grind pestle and tube SZ22, Kontes Glass Company, Vineland, New Jersey), each batch of 100 worms separately and all under a pure nitrogen atmosphere. *Ca*. Thiosymbion cells were separated from the host tissue and the other symbionts with a short centrifugation and snap frozen on liquid $N_2$.

Individual cell pellets were extracted with chloroform using the method described in detail elsewhere (Lemos et al). Briefly, cells were resuspended in 5 ml pure chloroform and possible PHAs were extracted by shaking the closed glass vials for 12h at 8°C. After centrifugation (10°C, 3000g, 5min) supernatants were transferred into new vials and dried under a constant stream of nitrogen. The residue was redissolved in 500 µL deuterochloroform ($CDCl_3$) containing trimethylsilane as internal standard. Samples were transferred into 7'inch NMR glass tubes. $^{1}$H-NMR and $^{13}$C-NMR spectra were obtained on a Bruker Avance DRX600 spectrometer with a 14.1 T magnet (Bruker Biospin; Rheinstetten, Germany) at 600.13 MHz and 140 MHz (300 K), respectively. For the 1D $^{1}$H spectra, a standard Bruker pulse sequence (NOESYpresat) was acquired over 12 kHz with a resolution of 32k data points; 64 scans were collected and summed per sample (Liebeke and Bundy 2013). $^{13}$C spectra were acquired using a standard 1D pulse sequence with decoupling, 1024 scans were acquired over a range of 200 kHz. Chemical shifts were referenced to the residual proton peak of $CDCl_3$ at 7.26 ppm and to the carbon peak of $CDCl_3$ at 77 ppm. Resulting peaks were matched with spectra from commercial available PHA mixtures and from literature values. Peaks for all detectable PHA forms (PHV, PHmV, PHB) were integrated and referenced again the internal standard for quantification. Amounts are given as total PHA based on the sum of signals belonging to the gamma position of each monomer.

##### *Optode measurements of single worm oxygen respiration*

Oxygen consumption rates of the holobiont were determined for freshly-collected white worms containing stored sulfur in *Ca*. Thiosymbion and for worms that had been incubated under oxic conditions prior to the measurements for 2 to 4 days to removed stored sulfur from *Ca*. Thiosymbion. The latter worms had subsequently changed their appearance from white to almost transparent (colourless pale). Sulfur content of all worms was determined after the respiration measurement as described above.

The oxygen respiration of single worms was determined by non-invasive measurement of oxygen concentration over time in a closed micro cuvette. Worms were incubated in a glass tube of approx. 40 µl volume containing sterile filtered seawater and about 10 glass beads (ø 0.4 - 0.6 mm, Sartorius, Göttingen), sealed with a butyl stopper. An oxygen optode (Holst & Grunwald 2001), which is a thin strip of transparent plastic foil coated with an oxygen-sensitive fluorescent polymer, was attached to the inside wall of the cuvette. During incubation the cuvette was slowly turned in a computerized rotor to mix the medium and stopped at intervals for the optical measurement of luminescence lifetime with the MOLLI system as described in Polerecky 2005. For each measurement the optode was excited by a flash from a blue light-emitting diode and the following fluorescence recorded with a CCD camera. The lifetime of the fluorescence depends on oxygen partial pressure at the optode and can thus be correlated to the oxygen concentration in the incubation cuvette. The respiration rate was calculated as the change of oxygen concentration over time at a range where the oxygen consumption was regarded linear (approx. 90 - 60 % oxygen saturation), corrected for the volume of the cuvette and normalized to the worm volume. To make the oxygen respiration rates comparable to the carbon bulk measurements volume normalized respiration rates were multiplied by the average biovolume of the worms used for carbon bulk analyses (0.532 µl = "average worm").

#### Modelling of carbon flow

##### *Model construction and assumptions*

We made the following assumptions for the model (Fig. 1):

1. Exchange of $CO_2$/bicarbonate between the seawater and the symbiosis is only limited by diffusion because the cuticle, which separates the seawater from the symbiosis is permeable for compounds up to 70 kDa in size (Dubilier et al., 2006).

2. Under the model conditions only the host and *Ca*. Thiosymbion are metabolically active and fix $CO_2$ for the following reasons: The sulfate-reducing δ-symbionts have been shown not to reduce sulfate under oxic conditions (Dubilier et al., 2001). The γ3-symbiont lacks the ability to use oxygen as $e^-$-acceptor relying on nitrate instead (Kleiner et al., 2012) and it also lacks an energy source, because it cannot store sulfur (Woyke et al., 2006). In *Ca*. Thiosymbion has internally stored sulfur as energy source and it can use oxygen as $e^-$-acceptor (Woyke et al., 2006; Kleiner et al., 2012).

3. Individual cells of one symbiont species are in different stages of their lifecycle and thus vary in size and carbon content. To account for this variability in carbon content between individual symbiont cells we implemented symbiont compartments as populations of individual cells with variable carbon contents. The average carbon content of symbiont cells can be calculated based on their known average biovolume (Ruehland et al., 2008) and their carbon to volume ratio (see section on Model Fitting below).

4. *Ca.* Thiosymbion cells divide into two equally sized daughter cells once they have doubled in size (e. g. TEM image in (Dubilier et al., 2001)).

5. Individual *Ca*. Thiosymbion cells have a variable amount of stored PHA and sulfur. When a *Ca*. Thiosymbion cell divides, each of the daughter cells will receive half of the remaining storage compounds.

6. *Ca.* Thiosymbion oxidizes sulfur at a cell specific rate ($r_s$) (Lenk et al., 2011) and uses the resulting energy to fix fresh carbon from the seawater DIC pool with a limited carbon yield stoichiometry (s2c). The parameter s2c was implemented as the theoretically maximal number of moles carbon fixed per mole sulfur, which based on the work of Kelly (1999) is 4.76 mol $CO_2$ fixed per mol of $S^0$ (Kelly, 1999). The s2c parameter specifies a percentage of this maximal carbon yield stoichiometry.

7. The worm, as all animals, fixes some $CO_2$ directly into its tissue by heterotrophic $CO_2$ fixation. Heterotrophic $CO_2$ fixation is ubiquitous in animals and results from anaplerotic refilling of the intermediary metabolism by the enzymes PEP carboxykinase and pyruvate carboxylase (Wood et al., 1941; Wood et al., 1945; Utter and Wood, 1946; Utter and Keech, 1960; Kresge et al., 2005). Both enzymes were found to be abundantly expressed by *O. algarvensis* (Kleiner et al., 2012). The amount of heterotrophic carbon fixation directly depends on metabolic activity and thus heterotrophic carbon fixation can be expressed as a fraction of respiration ($\varphi_h$) (Miltner et al., 2005).

8. *Ca*. Thiosymbion remobilizes the carbon stored in PHA with a rate ($\pi_{PHA}$) that is proportional to the amount of available PHA because with shrinking granule size the surface area to which the PHA depolymerase can bind (Pötter and Steinbüchel, 2006) gets limited.

9. Currently the fate of freshly fixed carbon and carbon from remobilized PHA in *Ca*. Thiosymbion is unknown. In theory this carbon could be used for growth by *Ca.* Thiosymbion ($\gamma_{PHA}$, $\gamma_f$) or be transferred to the host by excretion ($\varepsilon_{PHA}$, $\varepsilon_f$).

10. Carbon from all symbionts is acquired by the host through digestion of symbiont cells (Wippler et al., 2016). The digestion rate ($\pi_d$) is proportional to the size of the symbiont population.

11. Part of carbon assimilated by the host is used for growth ($\beta$ = production efficiency), whereas the rest (1-$\beta$) is used for respiratory energy gain leading to its release in form of $CO_2$ to the seawater DIC pool. For oligochaetes production efficiencies between 15 and 43% have been reported (Dash and Patra, 1977; Senapati and Dash, 1983; Mishra and Dash, 1984; Edwards and Bohlen, 1996; Jenderedjian, 1996).

12. Carbon is not respired by *Ca*. Thiosymbion because simultaneous autotrophic carbon fixation and heterotrophic carbon respiration would lead to futile cycling.

13. Transfer of carbon to the host by cell digestion or excretion is loss free (100% assimilation efficiency) i.e. all transferred carbon can actually be used by the host for growth and respiration.

To adapt our model to the conditions of our experiments (see above) we accounted for carbon that was already present in the symbiosis at the start of the experiment (mostly $^{12}C$) and freshly fixed carbon from the $^{13}C$-labeled DIC pool (Fig. 1). Additionally we included a parameter for label exchange with the atmosphere ($\tau_{ex}$) and considered the size of the seawater DIC pool in the incubations.

#### *Estimate parameter values by fitting with experimental data*

As the basis for fitting our model with the experimental data we defined an "average worm" based on the averages from our worm carbon bulk measurements. An average worm thus has a carbon content of 5 µmol and a biovolume of 0.532 µl. Wherever possible we used carbon content measures or biovolumes to convert rates and concentrations of different measurements to be comparable to the "average worm" (e. g. sulfur content and respiration rates).

Values for model parameters were determined as follows. For parameter values, which were directly measured in the experiments or could be derived by simple calculations from the experimental data, the mean value was entered into the model (Table S1). Then additional parameter values were estimated by fitting the model to specific experimental data (Table S1). For example, the symbiont digestion rate ($\pi_d$) was estimated using the respiration rate of an average low-sulfur worm (50 pmol $O_2$ $min^{-1}$, see results) and the assumption that in low-sulfur worms the only organic material that the worms have for respiration is derived from symbiont digestion because symbionts do not excrete carbon. For this estimation, $\pi_d$ was adjusted so that the modelled host respiration due to symbiont digestion equals 50 pmol $O_2$ $min^{-1}$. For some parameters direct measurement or indirect estimation from the experimental data was not possible. These parameters included the host production efficiency ($\beta$), host anaplerotic carbon fixation ($\varphi_h$), PHA remobilization rate ($\pi_{PHA}$), the fate of remobilized PHA ($\gamma_{PHA}$ and $\varepsilon_{PHA}$), and the fate of freshly fixed carbon ($\gamma_f$ and $\varepsilon_f$).Values for these parameters were estimated by finding a good fit of the model with all experimental data (e.g. $^{13}C$-abundances in *Ca.*

Thiosymbion cells and whole worms) while keeping all previously measured or estimated parameter values fixed and only varying the undetermined parameters.

##### *Sensitivity analyses*

Since the experimental data was not sufficient to unambiguously constrain all model parameters, in particular those describing physiological aspects of carbon transfer, the fitted model was used to explore the range of values that these parameters can realistically assume. For this, one or several of the unconstrained parameters were varied or co-varied and the model outcomes ($^{13}C$-abundances in different compartments, oxygen respiration rates and consumption of stored sulfur) were compared to the experimental data. A parameter value was considered to be “realistic” if the model outcomes stayed within a 95% confidence interval surrounding the means of important experimental data points (Table S1).

#### SI Results and Discussion

##### *Estimating the total biovolume of* Ca. *Thiosymbion in one worm*

We used the FISH images of the worm cross sections prepared for nanoSIMS analyses to estimate the volume occupied by *Ca*. Thiosymbion in the symbiosis. We compared the area of *Ca*. Thiosymbion (identified based on *Ca*. Thiosymbion probe signal) with the total area of the cross section. Areas were determined photogrammetrically with the software ImageJ (http://imagej.nih.gov/ij/) (Schneider et al., 2012). *Ca*. Thiosymbion accounted for 14 to 30% of the total area of worm cross sections with an average of 21% and thus the *Ca*. Thiosymbion population accounts for 0.074 to 0.16 µl (average 0.112 µl) of the biovolume of an average worm (0.532 µl).

##### *$^{13}C$-abundances in whole worms*

The natural $^{13}C$-abundance of worms at t=0 was 0.0107, which corresponds to a $\delta^{13}C$ value of -30.13‰ VPDB. As in the single cell analyses, significant $^{13}C$ enrichment in bulk analyses of whole worms was detected already after 10 minutes incubation ( $^{13}C/(^{13}C+^{12}C)$ = 0.0119, SE=0.0001, N=5 replicates, each with 3 pooled worms; ANOVA, p=0.0054)(Table S3). The $^{13}C$ abundance further increased after 25 hours ( $^{13}C/(^{13}C+^{12}C)$ = 0.0367, SE=0.0021, N=5), as in the single cell analyses, but the increase was on average only about 33% of that measured for individual *Ca*. Thiosymbion cells (Fig. 3b). The rate of increase in $^{13}C$ abundance decreased to about 35% of the initial rate in worms incubated for longer periods (8, 16 and 25 hours). The average carbon content of a whole worm was 5.0 µmol C $ind^{-1}$ (SE = 0.2, N=50) and its average biovolume was 0.532 µl (Table S3).

##### *Fraction of carbon in symbionts and symbiont numbers*

To estimate the fraction of whole symbiosis carbon that is contained in the symbionts, we used the symbionts known biovolumes and abundances (Ruehland et al., 2008), the experimentally determined carbon to volume ratio of *Ca*. Thiosymbion (see SI Results), and the experimentally determined amount of carbon contained in the *Ca*. Thiosymbion population (see SI Results). We assumed that the γ3-symbiont has the same high carbon to volume ratio as *Ca*. Thiosymbion (176.8 fg C $\mu m^{-3}$) because it also stores dense carbon in form of PHA in large quantities (Kleiner et al. unpublished data). For the remaining symbionts we used a carbon to volume ratio of 63 fg C $\mu m^{-3}$, which is the mean for bacteria as determined by Fagerbakke et al. (1996). Based on these data and assumptions the fraction of whole symbiosis carbon contained in the symbionts was 0.4.

##### *Changes in seawater $^{13}C$ abundance and estimation of $\tau_{ex}$*

The decrease of seawater $^{13}C$ abundance in incubations with worms was stronger as compared to the control incubations without worms. In the incubations with worms it decreased from 88 atom % $^{13}C$ at time point zero to 51 atom % $^{13}C$ after 25 hours, whereas in incubations without worms it decreased to 56 atom % $^{13}C$ after 25 hours.

The parameter $\tau_{ex}$, which characterizes the rate of $CO_2$ exchange with the atmosphere (Fig. 1), was estimated to be 61.8 hours by fitting the model to the seawater $^{13}C$ abundance values of the incubations without worms. For this purpose the worm and symbiont activities in the model were all set to zero.

##### *Estimating the carbon yield stoichiometry (s2c) in Ca. Thiosymbion*

We estimated s2c for *O. algarvensis* both by calculating it using our experimental data and by estimating the upper and lower limit using our model.

We calculated s2c using the average $S^0$ consumption in a worm within 25 hours (184 nmol) and the average $^{13}C$+$^{12}C$ incorporation (185.7 nmol). The average $^{13}C$+$^{12}C$ incorporation was calculated by correcting the average amount of $^{13}C$ incorporation derived from the bulk measurements (130 nmol) with the average fraction of $^{13}C$ in the seawater DIC pool (70%). The resulting fixed $CO_2$ to $S^0$ ratio was 1.01, which corresponds to 21.2% of the theoretical maximum of 4.76.

This apparent value for s2c can potentially differ from the actual carbon yield stoichiometry of *Ca*. Thiosymbion, because symbiosis internal factors that could not be directly included in the calculation may skew the apparent carbon yield stoichiometry. These factors include the heterotrophic $CO_2$ fixation by host tissue and the potential quick release of freshly fixed carbon to the seawater DIC pool through host respiration. To estimate the possible range for s2c, we used our model to estimate the lowest and highest possible values for it. For this we fitted the model to achieve the lowest and highest possible values for s2c, while staying within a reasonable range (95% confidence interval) of the averages of the experimentally determined values. The lowest possible value for s2c was 0.7 mol $CO_2$ per mol $S^0$ and the highest 2.2 mol $CO_2$ per mol $S^0$. Obtaining these minimal and maximal values required some extreme assumptions for the model parameters. To get to the lowest possible value for s2c the heterotrophic fixation by the host ($\varphi_h$) had to be 6% and host production efficiency ($\beta$) had to be 45%, both values are on the upper end of possible values when compared to the literature (SI methods). Additionally, the total amount of stored sulfur, sulfur consumption and initial oxygen respiration were at the upper limit of the 95% confidence interval surrounding the average of the experimental data, while initial PHA content was at the lower limit (Table S1). To get to the highest possible value for s2c, $\beta$ had to be 10%, $\varphi_h$ had to be 6% and a large share (31%) of freshly fixed carbon had to be directly transferred to the host by excretion ($\varepsilon_f$). Additionally, the total amount of stored sulfur, sulfur consumption and initial oxygen respiration were at the lower limit of the 95% confidence interval surrounding the average of the experimental data (Table S1).

#### *How do the carbon fixation rates of* **Ca.** *Thiosymbion compare to other autotrophs/SOX?*

The maximal gross carbon fixation rate of 360 amol $h^{-1}$ $cell^{-1}$ of *Ca.* Thiosymbion was comparable to those reported from other chemoautotrophic symbionts and around 28 fold lower than those of cultivable, free-living bacteria. For example, inorganic carbon uptake rates for the SOX symbiont of the fast-growing vent tubeworm *Riftia pachyptila* were 120 to 7,570 amol $h^{-1}$ $cell^{-1}$ (Fisher et al., 1989), for the SOX symbiont of the deep-sea mussel *Bathymodiolus thermophilus* 8–18 amol $h^{-1}$ $cell^{-1}$ (Belkin et al., 1986), and for the SOX symbiont of the shallow-water clam *Solemya velum* 1,300 to 7,580 amol $h^{-1}$ $cell^{-1}$ (Cavanaugh, 1983). For free-living SOX, much higher carbon uptake rates have been reported, for example, up to 10,000 amol $h^{-1}$ $cell^{-1}$ for the free-living purple sulfur bacterium *Chromatium okenii* (Musat et al., 2008).

The doubling time of 84 hours for *Ca.* Thiosymbion is in the same range as previously reported doubling times for other chemosynthetic symbionts. For example, the doubling time of the *B. thermophiles* symbiont was calculated to be between 60 to 140 hours (Belkin et al., 1986). As previously noted by Belkin et al. (1986) these doubling times are very long in comparison to free-living SOX isolated from hydrothermal vents, which can be less than one hour (Jannasch and Mottl, 1985).

#### *Quantification of energy efficiency of carbon fixation driven by sulfur oxidation in Ca. Thiosymbion based on Klatt and Polerecky (2015)*

Here we quantify the theoretical constraints on the energy efficiencies associated with the process of inorganic carbon fixation coupled to aerobic zero-valent sulfur ($S^0$) oxidation. Based on these we argue that *Ca*. Thiosymbion most likely uses another source of electrons in addition to $S^0$ for the reduction of $CO_2$ to organic carbon. Our calculations are based on basic principles of physical chemistry and employ the framework proposed by Klatt and Polerecky (2015) (Klatt and Polerecky, 2015).

##### Energy gaining vs. energy requiring processes

Energy required for $CO_2$ fixation is gained from the reduction of oxygen with $S^0$:

(Eq. 1a) <u>$O_2$ reduction:</u> $S^0 + 1.5\ O_2 + H_2O \rightarrow SO_4^{2-} + 2\ H^+$.

The Gibbs free energy of this reaction is

(Eq. 1b) $\Delta G_r(O_2\ red) = \Delta G_r^0(O_2\ red) + RT \ln Q_1$,

where

(Eq. 1c) $\Delta G_r^0(O_2\ red) = \Delta G_f^0(SO_4^{2-}) + 2\ \Delta G_f^0(H^+) - \Delta G_f^0(S^0) - 1.5\ \Delta G_f^0(O_2) - \Delta G_f^0(H_2O)$,

(Eq. 1d) $Q_1 = [SO_4^{2-}]\ [H^+]^2\ [O_2]^{-1.5}$,

$\Delta G_f^0$ refers to the standard Gibbs free energy of formation of the corresponding reactant, T is temperature (in K), and R is the gas constant. Note that the energy is expressed in kJ (mol $S^0)^{-1}$.

As argued by Klatt and Polerecky (2015), energy requirements for $CO_2$ fixation in sulfur oxidizing bacteria must be divided between two processes: the actual $CO_2$ reduction by NADH, and the production of the reducing equivalents NADH. For the Calvin cycle, $CO_2$ reduction occurs according to reaction

(Eq. 2a) <u>$CO_2$ reduction:</u> $3\ NADH + 1.5\ CO_2 + 0.5\ P_i \rightarrow 3\ NAD^+ + 0.5\ GA3P + 3.5\ H_2O$.

This reaction requires conversion of ATP into ADP and $P_i$, and its energy requirement is (see Supplemental material of Bar-Even et al., 2010 (Bar-Even et al., 2010))

(Eq. 2b) $\Delta G_r(CO_2\ red) = 69.7\ kJ\ (mol\ CO_2)^{-1}$.

With respect to the production of the reducing equivalents NADH, in this work we assume that the required electrons originate from *two sources*: $S^0$ and PHA, both internally stored in the gamma1 cells. In the first case, NADH is produced via membrane-associated reverse electron transport (RET) reactions with $S^0$ serving as the electron donor:

(Eq. 3a) <u>RET:</u> $S^0 + 3\ NAD^+ + 4\ H_2O \rightarrow SO_4^{2-} + 3\ NADH + 5\ H^+$.

The energy requirement of this reaction is, in kJ (mol $S^0$)$^{-1}$,

(Eq. 3b) $\Delta G_r(RET) = \Delta G_r^0(RET) + RT \ln Q_2$,

where

(Eq. 3c) $\Delta G_r^0(RET) = \Delta G_f^0(SO_4^{2-}) + 5\ \Delta G_f^0(H^+) - \Delta G_f^0(S^0) - 4\ \Delta G_f^0(H_2O) + 3\ [\Delta G_f^0(NADH) - \Delta G_f^0(NAD^+)]$,

(Eq. 3d) $Q_2 = [SO_4^{2-}]\ [H^+]^5\ [NADH]^3\ [NAD^+]^{-3}$,

and the difference $\Delta G_f^0(NADH) - \Delta G_f^0(NAD^+) = 60.99\ kJ\ (mol\ NADH)^{-1}$ (Alberty, 1998). In the second case, where NADH is produced by reducing $NAD^+$ using electrons from remobilized PHA, we assume that the energy requirement is *zero* (Martin et al., 2006).

**Overall efficiency**

To quantify the overall energy efficiency of $CO_2$ fixation coupled to the processes mentioned above, we make the following assumptions. First, we assume that for each mole of $S^0$, fraction *y* is used as the ultimate source of electrons for the reduction of $CO_2$ through reactions (2-3), while the remaining fraction 1-*y* is used to generate energy through reaction (1). Second, we assume that an extra *x* moles of $CO_2$ can be fixed through reaction (2) using NADH, and thus electrons, originating from other sources available to the cell, such as PHA.

Thus, by summing up equations (1) multiplied with 1-*y,* equations (2-3) multiplied with *y*, and equations (2) multiplied with *x*/1.5, we arrive at the following *net* reaction:

(Eq. 4) $S^0 + 1.5(1-y)\ O_2 + (1.5y+x)\ CO_2 + 2x\ NADH + (y/2+x/3)\ P_i + (1-y/2-7x/3)\ H_2O \rightarrow SO_4^{2-} + (2+3y)\ H^+ + (y/2+x/3)\ GA3P + 2x\ NAD^+$.

In other words, utilization of one mole of $S^0$ and 2*x* extra moles of NADH leads to the fixation of 1.5y+x moles of $CO_2$ and consumption of 1.5(1-y) moles of $O_2$. The energy required for these coupled processes is

$$\Delta G_r(required) = (1.5y+x)\ \Delta G_r(CO_2\ red) + y\ \Delta G_r(RET),$$

whereas the energy gained is

$$\Delta G_r(gained) = -\ (1-y)\ \Delta G_r(O_2\ red).$$

Therefore, the overall energy efficiency of the coupled process, defined as

$$e = \frac{\Delta G_r(\text{required})}{\Delta G_r(\text{gained})}$$

is calculated as

(Eq. 5) $$e = -\frac{(1.5y+x)\Delta G_r(CO_2\text{red})+y\Delta G_r(\text{RET})}{(1-y)\Delta G_r(O_2\text{red})}$$

Note that the efficiency depends on the parameters *x* and *y*, and thus on the stoichiometry of the net reaction (4). That is, provided that the stoichiometry of carbon fixation coupled to zero-valent sulfur oxidation is known, equation (5) makes it possible to *calculate* the overall energy efficiency of the process.

**Partial efficiencies**

In biochemical reactions, chemical energy gained from exergonic reactions is stored in ATP, while energy required for endergonic reactions is provided by ATP. Thus, it is convenient to associate with each reaction an efficiency that relates its true energy requirements with the energy requirements expressed in terms of ATP (Klatt and Polerecky, 2015). To do this, we denote the energy in the form of ATP that is stored or utilized during a particular reaction as $\Delta E_r$. Thus, we define the *efficiency of conversion of chemical energy to ATP associated with aerobic $S^0$ oxidation* (reaction 1) as

(Eq. 6) $e_{SO} = \Delta E_r(O_2 \text{ red}) / \Delta G_r^0(O_2 \text{ red})$.

Similarly, to relate the chemical energy stored in the form of organic carbon compounds (e.g., GA3P) to the energy in the form of ATP required to fix $CO_2$ (reaction 2), we define the *efficiency of the carbon fixation pathway* as

(Eq. 7) $e_{CO2} = \Delta G_r^0(CO_2 \text{ red}) / \Delta E_r(CO_2 \text{ red})$.

If we take the value of $\Delta G_r(CO_2 \text{ red}) = 69.7$ kJ (mol $CO_2)^{-1}$ calculated by Bar-Even et al. (2010)(Bar-Even et al., 2010), and assume that *Ca.* Thiosymbion cells require 3 ATP per $CO_2$ fixed, i.e., $\Delta E_r(CO_2 \text{ red}) = 123$ kJ (mol $CO_2)^{-1}$, we see that $e_{CO2} = 0.566$.

Finally, to express the chemical energy required for RET (reaction 3) in terms of ATP, we define the *efficiency of the RET reactions* as

(Eq. 8) $e_{RET} = \Delta G_r^0(\text{RET}) / \Delta E_r(\text{RET})$.

In addition to the efficiencies associated with the processes of energy conversion, we also define the *efficiency of ATP utilization for $CO_2$ fixation* as

(Eq. 9) $$e_u = -\frac{(1.5y+x)\Delta E_r(CO_2\text{red})+y\Delta E_r(\text{RET})}{(1-y)\Delta E_r(O_2\text{red})}.$$

Note that the expression for this efficiency is similar to that for the overall efficiency (Eq. 5), except for the fact that $\Delta E_r$ is used instead of $\Delta G_r$. As follows from this definition, $e_u$ represents the *fraction of ATP* gained from $S^0$ oxidation that is channeled towards and ultimately used for $CO_2$ fixation (i.e., both the $CO_2$ reduction and RET reactions).

Using these partial efficiencies, it is possible to rewrite the overall energy efficiency in Eq. 5 as

(Eq. 10) $e = (e_{SO}\ e_u) / f$,

where the factor f is defined as

(Eq. 11) $$f = \frac{1.5y\left(\frac{\alpha}{e_{CO2}} + \frac{1-\alpha}{e_{RET}}\right) + \frac{x\alpha}{e_{CO2}}}{1.5y + x\alpha},$$

and the parameter α is defined as

(Eq. 12) $$\alpha = \frac{1.5\Delta G_r(CO_2\text{red})}{1.5\Delta G_r(CO_2\text{red}) + \Delta G_r(\text{RET})}.$$

Note that unlike the factor f, the parameter α does not depend on the parameters *x* and *y*.

**Theoretical constraints on the partial efficiencies**

Now we apply the laws of thermodynamics to derive theoretical constraints on the partial efficiencies $e_{SO}$ and $e_{RET}$. Specifically, the energy provided by ATP for RET reactions must always be greater than or equal to the actual Gibbs free energy associated with the RET reactions. This implies $e_{RET} \leq 1$. Because $e_{CO2} = 0.566$ is fixed (see above), this constraint implies that

(Eq. 13) $$f \geq f_{min} = \frac{1.5y\left(\frac{\alpha}{e_{CO2}} + 1 - \alpha\right) + \frac{x\alpha}{e_{CO2}}}{1.5y + x\alpha}.$$

Additionally, assuming that $S^0$ oxidation is the only source of energy for *Ca*. Thiosymbion cells, the amount of ATP gained through $S^0$ oxidation must always be larger than or equal to the amount of ATP ultimately utilized for $CO_2$ fixation. This implies $e_u \leq 1$.

Combining these three constraints with Eq. (10), we derive the following constraint on the efficiency of $S^0$ oxidation in *Ca*. Thiosymbion cells:

(Eq. 14) $$e_{SO} \geq e_{SO,min} = e\, f_{min}.$$

This means that in order to satisfy laws of thermodynamics, the efficiency of chemical energy to ATP conversion during $S^0$ oxidation must always be at least $e_{SO,min}$.

Similar constraint can be derived for $e_{RET}$. Specifically, the amount of ATP generated from $S^0$ oxidation must always be lower than or equal to the Gibbs free energy released in this process. This implies $e_{SO} \leq 1$. Because $e_u \leq 1$, as already argued above, equation (10) implies $f \leq 1/e$. Combining this constraint with the definition of f (Eq. 11), we derive the following constraint on the efficiency of RET reactions in *Ca*. Thiosymbion cells:

(Eq. 15) $$e_{RET} \geq e_{RET,min} = \frac{1.5y(1-\alpha)}{\frac{1}{e}(1.5y + \alpha x) - \frac{\alpha}{e_{CO2}}(1.5y + x)}$$

This means that in order to satisfy laws of thermodynamics, the efficiency of ATP conversion to chemical energy during RET reactions must always be at least $e_{RET,min}$.

**Specific examples**

We implemented the above equations in R to calculate the constraints on the efficiencies $e_{SO}$ and $e_{RET}$ for a few specific conditions (R script available as Supplemental File S1). First, our experimental data indicate that $S^0$ consumption and $CO_2$ fixation in *Ca*. Thiosymbion cells occurs with approximately a 1:1 stoichiometry (see section on s2c). This implies that in the net equation (4) the stoichiometric coefficient $1.5y + x$ must be equal to 1. Because our measurements were insufficient to constrain the

$S^0$:$O_2$ or $CO_2$:$O_2$ stoichiometry, we were not able to simultaneously constrain both parameters $x$ and $y$. Therefore, we performed our calculations for a range of values of $x$ and $y$ such that $1.5y + x = 1$.

To calculate the overall efficiency from Eq. 5, we first substituted the tabulated values of the standard Gibbs free energies of formation (see, e.g., Thauer et al. 1977) to equations (1c) and (3c). When calculating the quotients $Q_1$ and $Q_2$ (Eqs. 1d and 3d) we assumed standard biochemical conditions (pressure 1 atm, temperature 25 ºC, reactant concentrations 1 M, pH = 7). This resulted in the values of the Gibbs free energies $\Delta G_r(O_2\ red) = -611.96$ kJ (mol $S^0)^{-1}$ and $\Delta G_r(RET) = 187.27$ kJ (mol $S^0)^{-1}$ (Eqs. 1b and 3b) and of the parameter $\alpha = 0.358$ (Eq. 12). Finally, we calculated the values of $f_{min}$ (Eq. 13), the overall efficiency e (Eq. 5), and the theoretical minimum efficiencies $e_{SO,min}$ (Eq. 14) and $e_{RET,min}$ (Eq. 15).

The results, which are summarized in Table S1, show that for the given $CO_2$:$S^0$ stoichiometry of 1:1, the value $x = 0$ would require that the partial efficiencies $e_{SO}$ and $e_{RET}$ be larger than 1. Since this would contradict laws of thermodynamics, our analysis therefore suggests that $x$ must be > 0, that is, at least some of the $CO_2$ fixed by the gamma1 cells must be reduced using an electron donor other than $S^0$. The critical boundary is $x = 0.08$, i.e., at least 8% of $CO_2$ must be fixed in this way. However, in this case the partial efficiencies $e_{SO}$ and $e_{RET}$ would have to be 1, i.e., the conversion of chemical energy to ATP during $S^0$ oxidation as well as the utilization of ATP for RET reactions would need to be 100% efficient, which is unrealistic. More realistic scenarios need to consider that between 20-100% of $CO_2$ are fixed without the need for $S^0$ to provide electrons. In this case the amount of $O_2$ utilized per mole of $S^0$ would increase from 0.7 to 1.5 moles, the overall energy conversion efficiency would decrease from 0.59 to 0.11, the theoretical minimum efficiency of $S^0$ oxidation would decrease from 0.78 to 0.2, and the theoretical minimum efficiency of the RET reactions would decrease from 0.61 to 0 (Table S5).

#### *Model predicts that most or all electrons from sulfur are used for $O_2$ reduction*

According to the equation for aerobic sulfur oxidation ($S^0 + 1.5\ O_2 + H_2O \rightarrow SO_4^{2-} + 2\ H^+$), 1.5 moles of $O_2$ are respired per mol of $S^0$ consumed. However, if electrons from sulfur are also used for $CO_2$ reduction the respired $O_2$ to consumed $S^0$ ratio (sox2oxy) is smaller than the maximum of 1.5. For example, when all electrons for $CO_2$ reduction are derived from $S^0$ and the ratio of $CO_2$ fixed to $S^0$ consumed (s2c) is 1 as determined above, sox2oxy will be 0.5 (Table S5) and in case half of the fixed $CO_2$ is reduced with electrons from sulfur then sox2oxy is 1.

We implemented a parameter for sox2oxy in the model and then tested with what sox2oxy values we could fit the measured oxygen respiration rates. A sox2oxy value of 1.5 provided the best fit for all experimental data (Table S1, Fig. 5). With a sox2oxy value of 1 we were still able to fit the experimental data, albeit with extreme assumptions, such as immediate transfer to the host and subsequent respiration of all remobilized PHA. A sox2oxy value of 0.5 could not be fitted with the data. This indicates that the majority or all of the electrons required for $CO_2$ reduction must come from a source other than $S^0$ (see Discussion).

#### *Why is the energy efficiency of sulfur oxidation based carbon fixation in Ca. Thiosymbion so high?*

The efficiency in *Ca.* T. algarvensis is unrealistically high for at least three reasons. First, factorization (i.e. analysis of the partial efficiencies that lead to the overall efficiency ‘e’) of ‘e’ showed that some partial efficiencies of ‘e’ e.g. for sulfur oxidation coupled to terminal electron acceptor reduction ($e_{SO}$) were higher than 100%, which is thermodynamically impossible, as this would imply that energy is generated in this energy requiring process (aka perpetual motion machine) (Table S5, SI results). This suggests that at least one additional factor is missing in the overall chemical reactions used to generate the equations (SI Results). One potential factor that would influence the outcome of the calculations is that the CBB cycle in *Ca.* Thiosymbion might use less ATP compared to the textbook version of this pathway. Indeed, it was recently predicted that the CBB cycle in *Ca.* Thiosymbion may use as little as 2 ATP per $CO_2$ fixed as compared to 3 ATP in the textbook version (Kleiner et al., 2012b). We recalculated the energy conservation efficiency using this lower ATP requirement (SI Results). The overall efficiency ‘e’ did not change (95%), but the partial efficiencies changed. The $CO_2$ fixation efficiency $e_{CO2}$ increased from 56.6% to 85%, whereas all other partial efficiencies were close to 100% (SI Results), which would suggest that sulfur oxidation, reverse electron transport and energy transfer are ~100% efficient. Since efficiencies of 100% violate the second law of thermodynamics, which states that in any thermodynamic process some energy is transformed into thermal energy and thus, in a biological system, effectively lost, this result cannot be accurate and we need to consider at least one additional factor. The second reason why the efficiency in *Ca.* T. algarvensis is unrealistically high, is that it has been shown that the key enzyme of the CBB cycle, the ribulose bisphosphate carboxylase/oxygenase (RubisCO) is inefficient, it is a bifunctional enzyme that catalyzes both the carboxylation and oxygenation of ribulose-1,5-bisphosphate. When the oxygenation reaction occurs energy for carbon fixation is effectively lost. Research on plants has shown that losses due to this so called photorespiration can be substantial in the range of 21 to 50% of all $CO_2$ assimilated (Peterson, 1983; Sharkey, 1988; Cegelski and Schaefer, 2006). Third, overall efficiency of energy transfer to the CBB cycle should be reduced due to energy consuming basic cell maintenance processes such as nutrient transport, DNA repair and removal of denatured proteins and biosynthesis steps, which convert hexoses produced by the CBB cycle into “permanent” biomass such as cell wall components, DNA and proteins.

### SI Figures

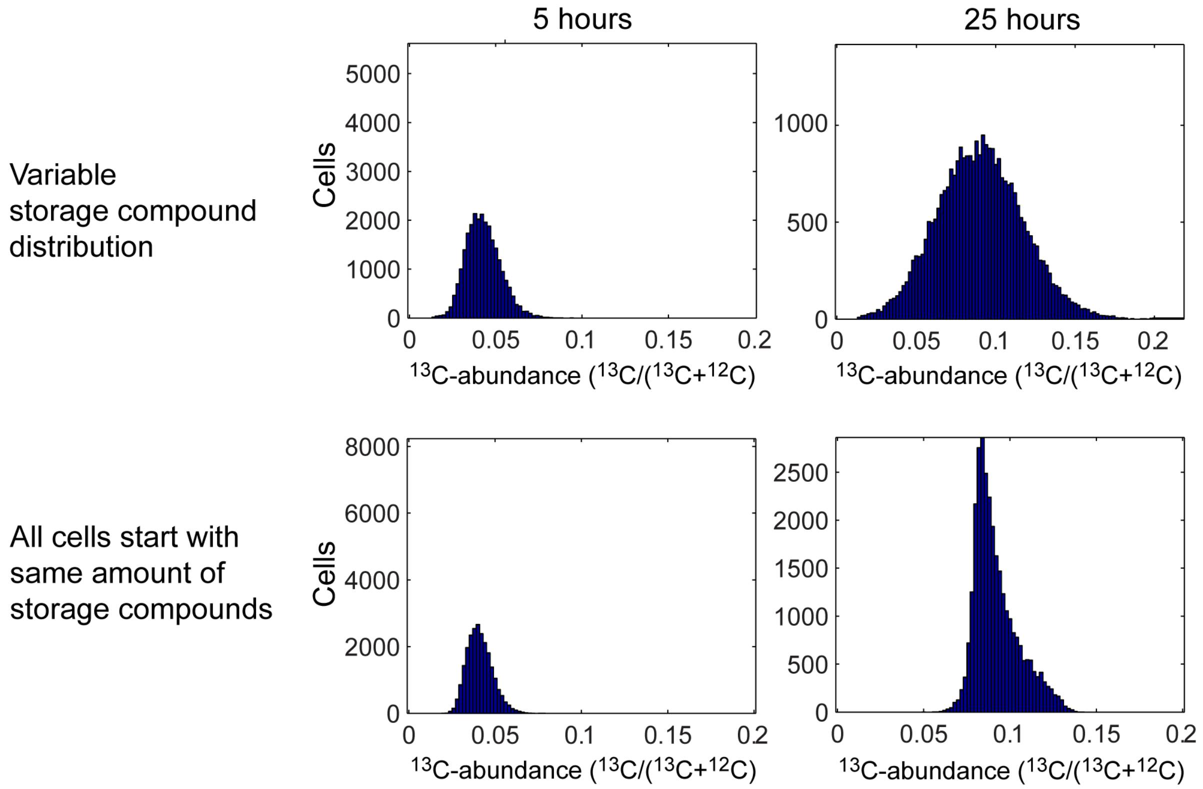

**Figure S1**: Modelled $^{13}$C-enrichments of individual *Ca*. Thiosymbion cells at 5 and 25 hours of oxic incubation. The top row shows the distribution of $^{13}$C-enrichment if cells start out with variable storage compound content. The bottom row shows the distribution of $^{13}$C-enrichment if cells all start out with the same exact amount of storage compounds. In each plot values for 20,000 cells are presented. Please note, the axes scaling is variable.

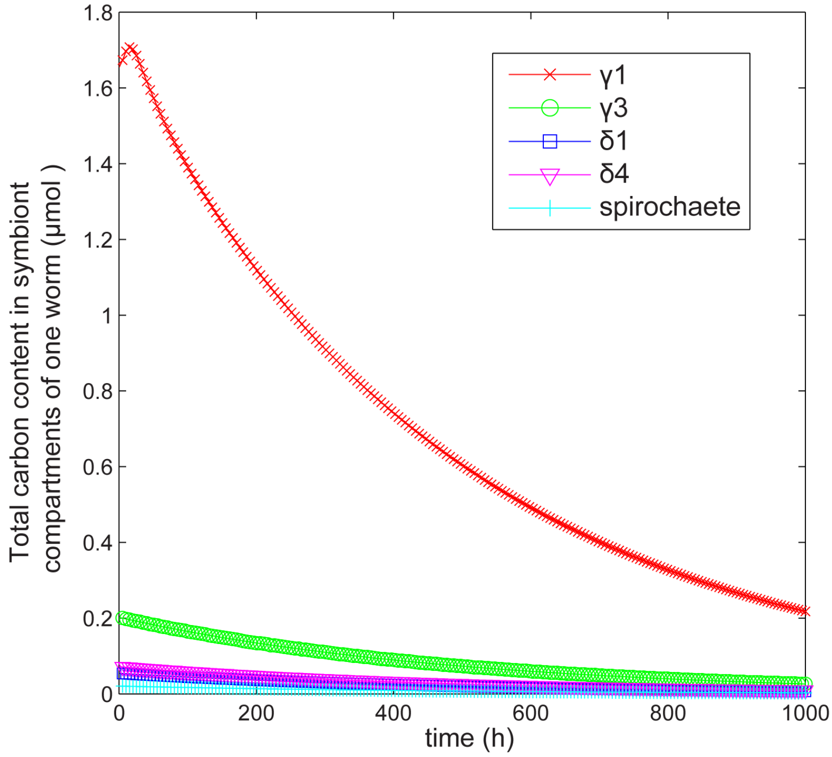

**Figure S2:** Modelled carbon pool of the symbiont population in one worm over time. The model was used to extrapolate the carbon pool changes up to 1,000 hours of oxic conditions.

### SI Tables

#### **Table S1:** Overview of experimental data used for model fitting and sensitivity analyses

For the sensitivity analyses 95% confidence intervals of means were used to determine upper and lower bounds of modelled means.

| **Parameter (Unit)** | **Mean** | **95% confidence interval of mean lower boundary** | **95% confidence interval of mean upper boundary** |
|---|---|---|---|
| $^{13}$C-abundance of *Ca*. Thiosymbion cells at 25 h based on NanoSIMS ($^{13}$C/($^{13}$C+$^{12}$C)) | 0.0898 | 0.0837 | 0.0960 |
| $^{13}$C-abundance in host tissue at 25 h based on NanoSIMS ($^{13}$C/($^{13}$C+$^{12}$C)) | 0.0130 | 0.0124 | 0.0136 |
| $^{13}$C-abundance of whole worms at 25 h ($^{13}$C/($^{13}$C+$^{12}$C)) | 0.0367 | 0.0325 | 0.0408 |
| Sulfur content of average worm at 0 h (nmol $S^0$) | 212.7 | 170.4 | 255.0 |
| Sulfur content of average worm at 25 h (nmol $S^0$) | 28.7 | 9.6 | 47.9 |
| Sulfur consumption of average worm within 25 h (nmol $S^0$) | 184.0 | 122.5 | 245.4 |
| Oxygen respiration low-sulfur worm (pmol $O_2$ /min*worm) | 50.11 | 34.83 | 65.39 |
| Oxygen respiration high-sulfur worm (pmol $O_2$ /min*worm) | 444.62 | 338.93 | 550.31 |

| | | | |
|---|---|---|---|
| $^{13}$C-abundance in seawater DIC at 25 h ($^{13}$C/($^{13}$C+$^{12}$C)) | 0.5114036 | 0.5096 | 0.5132 |
| Carbon in PHA/total carbon in *Ca*. Thiosymbion | 0.42 | 0.35 | 0.55 |

#### Table S2: Elemental sulfur content of worms from incubation experiment

Sulfur contents of individual worms were normalized to their biovolume and averages per time point were then multiplied by the average biovolume of the worms used for the $^{13}$C bulk measurements

| time point | N | nmol $S^0$ / average worm | STDEV | 95% confidence interval of mean upper boundary | 95% confidence interval of mean lower boundary |
|---|---|---|---|---|---|
| 0 min | 20 | 212.7 | 96.5 | 255.0 | 170.4 |
| 10 min | 12 | 205.7 | 103.5 | 264.3 | 147.1 |
| 20 min | 12 | 249.2 | 90.1 | 300.1 | 198.2 |
| 30 min | 13 | 159.5 | 89.4 | 208.1 | 110.8 |
| 1 h | 13 | 132.3 | 121.5 | 198.3 | 66.2 |
| 2 h | 12 | 134.9 | 130.1 | 208.6 | 61.3 |
| 4 h | 17 | 260.7 | 165.1 | 339.1 | 182.2 |
| 6 h | 19 | 207.1 | 119.7 | 261.0 | 153.3 |
| 8 h | 20 | 153.4 | 100.9 | 197.6 | 109.2 |
| 16 h | 19 | 158.7 | 108.5 | 207.5 | 110.0 |
| 25 h | 8 | 28.7 | 27.6 | 47.9 | 9.6 |

#### **Table S3:** Overview of worm bulk measurements

Five replicates containing three worms each were measured per time point

| TimePoint | Average $^{13}$C-content | STDEV $^{13}$C-content | 95% confidence interval of mean upper boundary | 95% confiden ce interval of mean lower boundar y | Avera ge µmol C per worm |
|---|---|---|---|---|---|
| 0 min [1] | 0.0107 | 0.0000 | 0.0107 | 0.0107 | 2.55 |
| 10 min | 0.0119 | 0.0003 | 0.0121 | 0.0116 | 3.77 |
| 20 min | 0.0120 | 0.0002 | 0.0121 | 0.0119 | 5.12 |
| 30 min | 0.0122 | 0.0004 | 0.0125 | 0.0119 | 4.78 |
| 60 min | 0.0136 | 0.0003 | 0.0139 | 0.0134 | 6.07 |
| 2 h | 0.0159 | 0.0012 | 0.0170 | 0.0149 | 4.36 |
| 4 h | 0.0194 | 0.0015 | 0.0207 | 0.0181 | 5.09 |
| 6 h | 0.0204 | 0.0016 | 0.0219 | 0.0190 | 5.06 |
| 8 h | 0.0213 | 0.0020 | 0.0231 | 0.0195 | 5.26 |
| 16 h [2] | 0.0267 | 0.0037 | 0.0304 | 0.0230 | 5.32 |
| 25 h | 0.0367 | 0.0048 | 0.0408 | 0.0325 | 5.08 |

**1**: Only three replicates measured for t = 0 min

**2**: Only four replicates measured for t = 16 h

**Table S4:** Overview of worm respiration rates

**Low-sulfur worms (elemental sulfur content 5.5 ± 6 nmol $mm^{-3}$)**

| Worm | Rate [nmol $O_2$/min] | Rate [pmol $O_2$/min] | Worm volume [mm3] | Volume normalized rate [pmol $O_2$ /min *$mm^3$] | Rate normalized to average bulk worm of 0.532 $mm^3$ volume [pmol $O_2$ /min*worm] |
|---|---|---|---|---|---|
| 1 | 0.0233 | 23.28 | 0.17 | 137.74 | 73.28 |
| 2 | 0.0342 | 34.22 | 0.60 | 56.66 | 30.14 |
| 3 | 0.0309 | 30.93 | 0.27 | 114.12 | 60.71 |
| 4 | 0.0252 | 25.16 | 0.24 | 105.29 | 56.01 |
| 5 | 0.0208 | 20.78 | 0.47 | 44.59 | 23.72 |
| 6 | 0.0479 | 47.92 | 0.45 | 106.73 | 56.78 |
| Average | | 30.38 | | 94.19 | 50.11 |
| STDEV | | 9.93 | | 35.90 | 19.10 |
| 95% confidence interval of mean upper boundary | | | | | 65.39 |
| 95% confidence interval of mean lower boundary | | | | | 34.83 |

**High-sulfur worms (elemental sulfur content 506 ± 103 nmol $mm^{-3}$)**

| Worm | Rate [nmol $O_2$/min] | Rate [pmol $O_2$/min] | Worm volume [mm3] | Volume normalized rate [pmol $O_2$ /min *$mm^3$] | Rate normalized to average bulk worm of 0.532 $mm^3$ volume [pmol $O_2$ /min*worm] |
|---|---|---|---|---|---|
| 7 | 0.3137 | 313.73 | 0.32 | 971.31 | 516.74 |
| 8 | 0.3825 | 382.53 | 0.69 | 556.81 | 296.22 |
| 9 | 0.4420 | 441.98 | 0.56 | 785.05 | 417.65 |
| 10 | 0.4020 | 401.97 | 0.32 | 1256.15 | 668.27 |
| 11 | 0.3497 | 349.67 | 0.45 | 782.25 | 416.16 |
| 12 | 0.4183 | 418.30 | 0.63 | 662.92 | 352.67 |
| Average | | 384.70 | | 835.75 | 444.62 |
| STDEV | | 46.86 | | 248.28 | 132.09 |
| 95% confidence interval of mean upper boundary | | | | | 550.31 |
| 95% confidence interval of mean lower boundary | | | | | 338.93 |

**Table S5: Theoretical constraints on the efficiencies of processes involved in aerobic $S^0$ oxidation coupled to $CO_2$ fixation as a function of stoichiometry based on Klatt and Polerecky (2015) (Klatt and Polerecky, 2015)**

| x | y | $CO_2$:$S^0$ = 1.5*y+x | $O_2/S^0$ = 1.5*(1-y) | $f_{min}$ | e | $e_{SO,min}$ | $e_{RET,min}$ |
|---|---|---|---|---|---|---|---|
| 0 | 0.666 | 1 | 0.5 | 1.27 | 0.953 | 1.22 | 1.54 |
| 0.08 | 0.613 | 1 | 0.58 | 1.29 | 0.78 | 1.00 | 1.01 |
| 0.1 | 0.6 | 1 | 0.6 | 1.29 | 0.74 | 0.96 | 0.92 |
| 0.2 | 0.533 | 1 | 0.7 | 1.31 | 0.59 | 0.78 | 0.61 |

| | | | | | | | |
|---|---|---|---|---|---|---|---|
| 0.3 | 0.466 | 1 | 0.8 | 1.34 | 0.48 | 0.64 | 0.43 |
| 0.4 | 0.4 | 1 | 0.9 | 1.37 | 0.39 | 0.54 | 0.31 |
| 0.5 | 0.333 | 1 | 1 | 1.40 | 0.32 | 0.45 | 0.22 |
| 0.6 | 0.266 | 1 | 1.1 | 1.45 | 0.26 | 0.39 | 0.15 |
| 1 | 0 | 1 | 1.5 | 1.76 | 0.11 | 0.2 | 0 |

x: Moles of $CO_2$ fixed using reducing equivalents not derived from $S^0$.
y: Fraction of $S^0$ used as source of electrons for the reduction of $CO_2$. 1-y is used for energy generation.
e: Overall energy efficiency of the coupled energy gaining and requiring processes.
$e_{S0}$: Partial efficiency of conversion of chemical energy to ATP associated with aerobic $S^0$ oxidation.
$e_{RET,min}$: Partial efficiency of the reverse electron transport (RET) reaction $e_{RET}$, must be larger or equal to $e_{RET,min}$ to satisfy the laws of thermodynamics.

**Table S6:** Number of worms per incubation dish

| incubation time | 10‘ | 20‘ | 30‘ | 1h | 2h | 4h | 6h | 8h | 16h | 25h |
|---|---|---|---|---|---|---|---|---|---|---|
| number of worms | 34 | 33 | 34 | 34 | 35 | 39 | 41 | 41 | 40 | 29 |

### SI Datasets

Submitted as separate files.

1. Zip folder containing the model files in matlab format
2. Intitial conditions file for model (annotated)
3. R-script for efficiency calculations (File S1)
4. Model output with final parameters (File S2)
